# A key GPCR phosphorylation motif discovered in arrestin2•CCR5 phosphopeptide complexes

**DOI:** 10.1101/2022.10.10.511578

**Authors:** Polina Isaikina, Ivana Petrovic, Roman P. Jakob, Parishmita Sarma, Ashutosh Ranjan, Minakshi Baruah, Vineet Panwalkar, Timm Maier, Arun K. Shukla, Stephan Grzesiek

**Affiliations:** Focal Area Structural Biology and Biophysics, Biozentrum, University of Basel, CH-4056 Basel, Switzerland; Department of Biological Sciences and Bioengineering, Indian Institute of Technology, Kanpur, India

**Keywords:** beta-arrestin, arrestin, G protein-coupled receptor (GPCR), CCR5, chemokine, phosphorylation, phosphopeptide, X-ray crystallography, NMR

## Abstract

The two non-visual arrestin isoforms, arrestin2 and arrestin3 recognize and bind hundreds of G protein-coupled receptors (GPCRs) with different phosphorylation patterns leading to distinct functional outcomes. The impact of phosphorylation on arrestin interactions has been well studied only for very few GPCRs. Here we have characterized the interactions between the phosphorylated CC chemokine receptor 5 (CCR5) and arrestin2. We detected several new CCR5 phosphorylation sites, which are necessary for stable complex formation with arrestin2. Crystal structures of arrestin2 in apo form and in complexes with CCR5 C-terminal phosphopeptides together with NMR spectroscopy, biochemical and functional assays revealed three phosphoresidues in a pXpp motif that are essential for the arrestin2 interactions and activation. The same phosphoresidue cluster is present in other receptors, which form stable complexes with arrestin2. We propose that the identified pXpp motif is responsible for robust arrestin2 recruitment in many GPCRs. An analysis of available sequences, structural and functional information on other GPCR•arrestin interactions suggests that a particular arrangement of phosphoresidues within the GPCR intracellular loop 3 and C-terminal tail determines arrestin2 and 3 isoform specificity. Taken together, our findings demonstrate how multi-site phosphorylation controls GPCR•arrestin interactions and provide a framework to probe the intricate details of arrestin activation and signaling.

**One-sentence summary:** A structural and functional analysis of arrestin2 in apo form and complexes with several CCR5 phosphopeptides reveals key phosphorylation sites responsible for stable GPCR•arrestin interactions and their contributions to CCR5-arrestin2 function.

## Introduction

GPCRs represent a large family of cell-surface receptors mediating signaling events via G proteins and arrestins^1^. The agonist-induced G protein signaling is terminated by the phosphorylation of the receptor C-terminal tail and/or intracellular loops^2^ primarily via GPCR kinases (GRKs), which subsequently leads to the binding of arrestins to the receptor C-terminal tail and core^3–6^. The receptor•arrestin complex acts as a scaffold for various further signaling proteins thereby activating e.g. ERK1/2 and MAP or inducing receptor internalization^2,7^. The many hundreds non-visual GPCRs in the human body are regulated by two genetically and structurally conserved non-visual arrestin subtypes, arrestin2 and arrestin3 (also known as β-arrestin1 and β-arrestin2, respectively). While both isoforms can bind to the same receptor, their interaction may activate different signaling partners^8^. The arrestin-mediated signaling depends on the phosphorylation pattern of the GPCR C-terminal tail, which is modulated by the interactions of the GPCR with various GRKs^9,10^. These findings have led to the hypothesis of a phosphorylation ‘barcode’ for the receptor C-terminal tail^11^.

The molecular details of the specificity of GPCR-arrestin interactions and the causes for the different functional outcomes of varying phosphorylation patterns are still poorly understood. Previously solved full-length GPCR•arrestin complex structures comprise a rhodopsin-arrestin1 fusion complex^12^, a fusion complex of the engineered constitutively active arrestin2 with the truncated 5HT_2B_ serotonin receptor^13^, an arrestin2 complex with a chimera of the M2 muscarinic receptor (M2R) and the vasopressin 2 receptor C-terminal phosphopeptide (V2Rpp)^14^, a β1AR-V2Rpp chimera•arrestin2 complex^15^, as well as native V2R•arrestin2^16^ and NTR1•arrestin2^17,18^ complexes. Of these, the rhodopsin fusion, the M2R-V2Rpp and β1AR-V2Rpp chimera, as well as the truncated 5HT_2B_ serotonin receptor fusion complexes show similar orientations of the arrestin. However, the arrestin2 orientation differs significantly in the NTR1 and V2R complexes, which have native receptor C-terminal tails. The phosphate groups in the C-terminal tails have only been resolved in the M2R-V2Rpp, β1AR-V2Rpp, and V2R complexes. Higher resolution has been obtained in complex structures of arrestins with phosphorylated GPCR C-terminal tail peptides, which have revealed the position and coordination of several phosphates in addition to the hallmarks of arrestin activation, i.e. the replacement of arrestin strand 20 by the phosphorylated receptor peptide, conformational changes within the arrestin loops and a twist between its N- and C-terminal domains. However, these studies have been limited to V2Rpps binding arrestin2^19,20^, an ACKR3 (formerly known as CXCR7) C-terminal phosphopeptide binding arrestin3^21^, and a rhodopsin C-terminal phosphopeptide binding arrestin1 (visual arrestin)^22^.

We have recently determined the structure of the human chemokine receptor 5 (CCR5) in an active complex with the chemokine super-agonist [6P4]CCL5 and the heterotrimeric G_i_ protein by cryo electron microscopy^23^. CCR5 plays a major role in inflammation by recruiting and activating leukocytes^24^. It is also the principal HIV coreceptor^25^ and is involved in the pathology of cancer^26,27^, neuroinflammation^24^, and COVID-19^28^. The structure of the active [6P4]CCL5•CCR5•G_i_ complex has provided detailed insights into the mechanism of CCR5 activation via the [6P4]CCL5 N-terminus, which differs from other solved chemokine•GPCR complexes.

Much less is known about the molecular basis of chemokine receptor interactions with arrestins, which play a similar critical role as G protein interactions in the immune response and inflammatory signaling pathways^29^. Here we have characterized the CCR5 phosphorylation induced by the super-agonist [6P4]CCL5 and GRK2. We discovered several new phosphorylation sites within the CCR5 C-terminal tail and used them together with previously reported sites to design a set of phosphopeptides for analyzing their effects on arrestin interactions. We determined high-resolution crystal structures of arrestin2 in apo form and in complex with two of these phosphopeptides with resolved positions of the phosphate groups in their electron density. The phosphate groups form a distinct pXpp phosphopeptide sequence motif that induces the active arrestin2 conformation by dominant electrostatic and beta-sheet interactions. An identical pXpp motif exists in the V2Rpp and forms the same activating interactions with arrestin2. A combination of NMR, biochemical and cellular assays confirmed the importance of this motif and quantified the contributions of individual phosphoresidues to arrestin binding and activation. A comparison of the new structures to other arrestin complexes together with an analysis of GPCR sequences provides hints on the molecular basis of arrestin2/arrestin3 isoform specificity.

## Results

### Revisiting CCR5 phosphorylation by a G protein kinase

Agonist-driven phosphorylation of serine and threonine residues in GPCRs by GRKs is generally required for arrestin binding. Although CCR5 has seven such potential phosphorylation sites in its C-terminal tail, phosphorylation of only four C-terminal serine residues has been reported ^30,31^. To clarify this situation, we characterized CCR5 phosphorylation by GRK2 that plays a prominent role in the regulation of many chemokine receptors, including CCR5 ^32–35^. For this, GRK2 was co-expressed with CCR5 in insect cells and phosphorylation induced by the addition of the super-agonist chemokine [6P4]CCL5. Phosphoproteomics of the purified CCR5 clearly revealed phosphorylation of S349 and S325, which had not previously been reported, in addition to a double phosphorylation of S336 and S337, as well as a not well-defined single phosphorylation site in the region T340 to T343 (Figures 1A, S1). Similar assignment ambiguities have been related previously to the heterogeneity of phosphorylation by GRKs ^18^. A further western blot analysis with CCR5 phosphosite-specific antibodies confirmed the previously not reported phosphorylation of T340 as well as that of the serine residues S336/337, S342, and S349 (Figure 1B).

**Figure 1.**
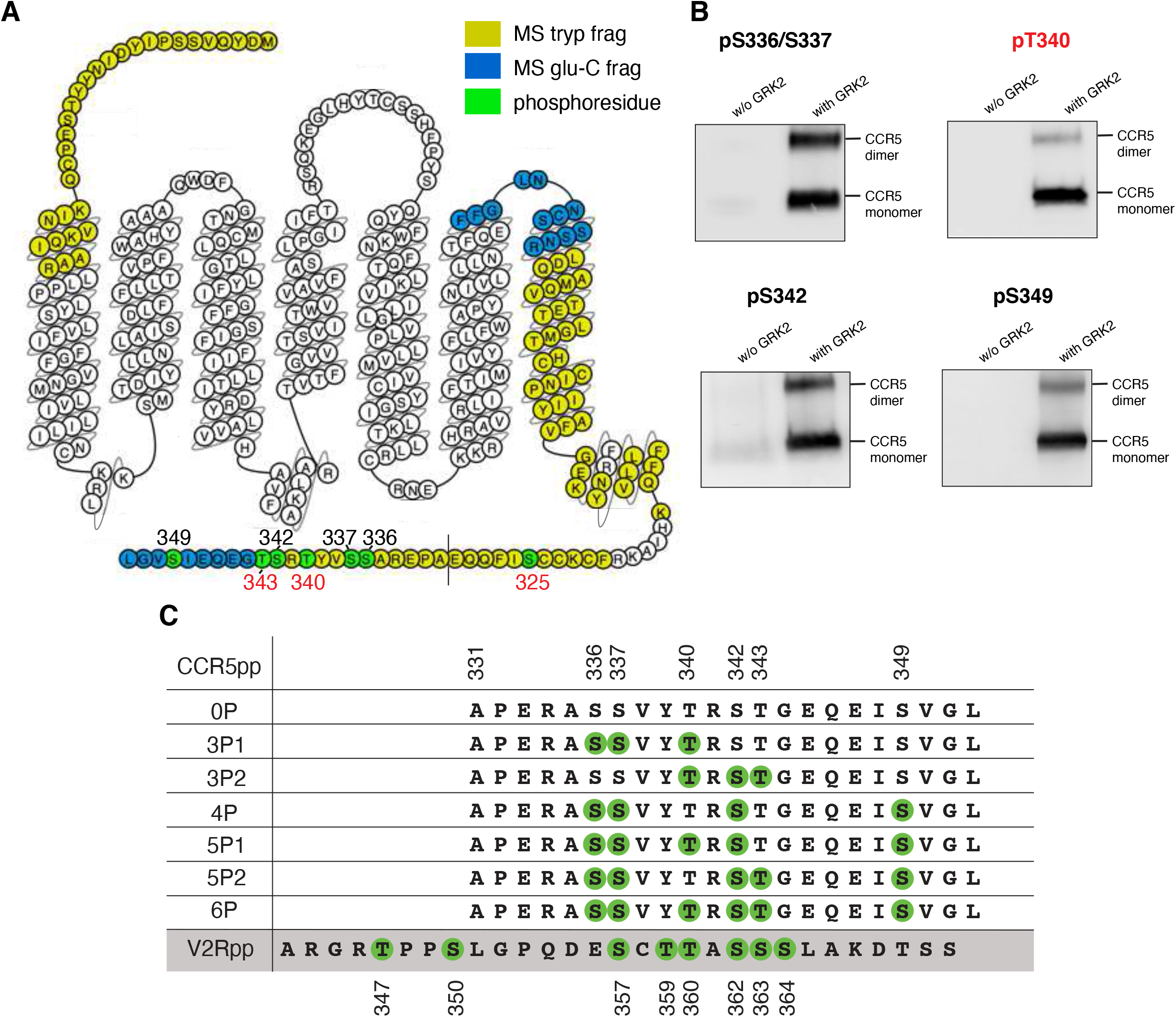
Detection of CCR5 phosphorylation and synthetic phosphopeptide design. (A) Mass spectrometric sequence coverage of GRK2-phosphorylated CCR5 using trypsin (yellow) and glu-C digestion (blue). Phosphosites detected by mass spectrometry are indicated in green. (B) Confirmation of mass spectrometry phosphorylation by western blot analysis with phosphosite-specific antibodies. (C) Designed synthetic phosphopeptides (0P–6P) corresponding to the last 22 residues of CCR5 based on the detected CCR5 phosphorylation. A further V2R C-terminal phosphopeptide (V2Rpp) was used as a control. Serine or threonine phosphorylation are indicated by a green circle.

To assess the impact of these various CCR5 phosphoresidues on arrestin2 binding and conformation, we designed seven distinct CCR5 synthetic phosphopeptides in addition to the V2Rpp phosphopeptide mimicking V2R phosphorylation ^20^, which served as a control (Figure 1C). These designed CCR5 phosphopeptides did not include pS325, which is unlikely to play a major role in the arrestin interaction due to its location at the beginning of the CCR5 C-terminal tail directly following the palmitoylated cysteine residues (Figure 1A). This is confirmed by functional assays (see below).

### CCR5 phosphorylation levels govern arrestin2 binding

The interaction of the synthesized phosphopeptides with arrestin2 was first investigated by solution NMR spectroscopy using a truncated arrestin construct (arrestin2^1-393^, residues 1-393), which comprises the C-terminal strand β20, but lacks the disordered C-terminal tail. Upon addition of the phosphopeptides, the ^1^H-^15^N HSQC-TROSY spectra of arrestin2^1-393^ [^15^N-labeled, ~80% deuterated] show continuous shifts in several resonance positions (Figure 2A, S2) indicating fast to intermediate chemical exchange on the microsecond time scale. A non-linear fit of the chemical shift changes to binding isotherms provided dissociation constants K_D_ in the ten to hundred micromolar range (Figures 2A, S2), which is similar to previous observations on phosphopeptide•arrestin1 complexes ^22^.

**Figure 2.**
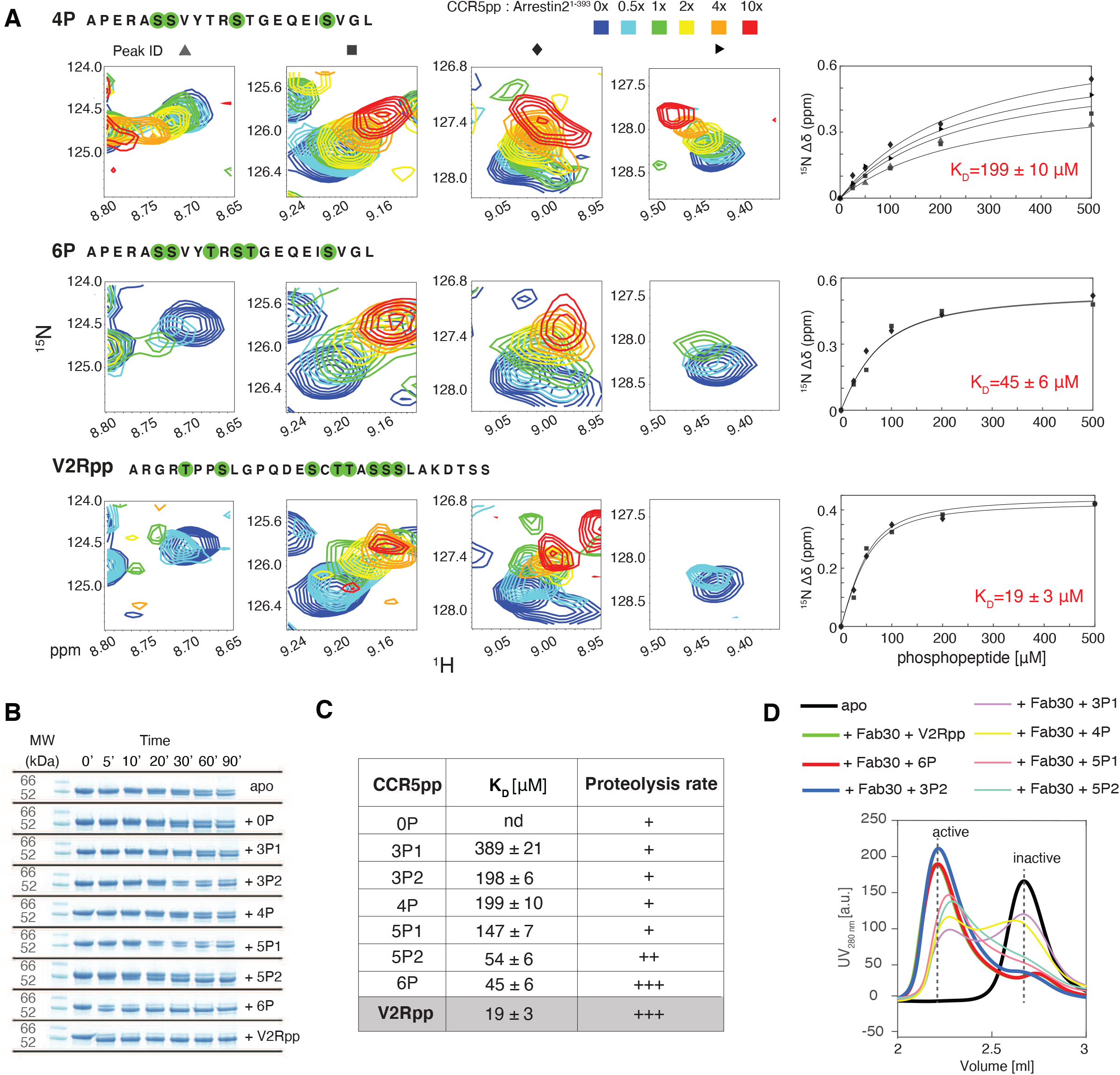
Arrestin2 interaction with CCR5 phosphopeptides. (A) Left: small regions of ^1^H-_15_N TROSY spectra showing resonance shifts of four selected arrestin2^1-393^ residues upon CCR5 4P, 6P and V2Rpp binding. Right: detected chemical shift changes as a function of phosphopeptide concentration. Solid lines depict global non-linear least-square fits to the data points with respective dissociation constants (see Methods). Analogous fits for the other phosphopeptide titrations are shown in Figure S1. (B) Trypsin proteolysis assay of arrestin2^1-418^ in apo form and in complexes with various phosphopeptides visualized by SDS-PAGE. (C) Summarized NMR titration and trypsin proteolysis results. Tighter peptide binding is correlated with faster arrestin2 digestion. (D) SEC profiles showing arrestin2 activation in the presence of phosphopeptides and Fab30. The active arrestin2•phosphopeptide complexes, which are recognized by Fab30, elute at lower volumes than the inactive apo form. Arrestin2 in presence of CCR5 6P, 3P2 and V2Rpp elutes only as active complexes, whereas the other peptides lead to mixtures of inactive and active arrestin2 forms.

Functional binding of the phosphopeptides to arrestin is expected to induce a conformational change, where arrestin’s C-terminal strand β20 is released from the β-sheet with its N-terminal β-strand and replaced by the phosphopeptide (see below). This exposes previously inaccessible cleavage sites in the arrestin C-terminus (R393 for arrestin2), which can be probed by trypsin digestion ^36^. Such a trypsin proteolysis carried out on full-length arrestin2 (arrestin2^1-418^, Figure 2B) confirmed the functional conformational changes induced by the binding of the synthetic phosphopeptides.

Figure 2C summarizes the quantitative results of the NMR titrations and the trypsin proteolysis assays. The tightest binding was observed for the six-fold phosphorylated CCR5 6P peptide (K_D_ = 45 ± 6 μM) and the eight-fold phosphorylated V2Rpp (K_D_ = 19 ± 3 μM), whereas the three- and four-fold phosphorylated peptides have weaker affinities in the hundred micromolar K_D_ range, agreeing with the expectation that the phosphorylation level dominates the arrestin•peptide interaction. However, individual phosphorylation sites contribute differently to the overall affinity. Hence, 5P1 (lacking pT343) shows weaker affinity (K_D_ = 147 ± 7 μM) whereas 5P2 (comprising pT343) has an affinity (K_D_ = 54 ± 6 μM) similar to 6P suggesting that phosphorylation of T343 is essential for tight arrestin binding.

The trypsin proteolysis rates of the arrestin•phosphopeptide complexes correlate with the determined affinities. Whereas the low-affinity (K_D_ > 100 μM) peptides (5P1 and all three- or four-fold phosphorylated peptides) had a trypsin digestion midpoint at around 30 minutes, for both 6P and V2Rpp the digested band became dominant already after only 5 minutes (Figure 2B). Of note, 5P2 comprising pT343 also had an accelerated digestion midpoint at about 20 minutes.

To probe active arrestin2 conformation induced by the synthetic CCR5 phosphopeptides, we incubated full-length arrestin2 with the peptides and synthetic antibody fragment Fab30 that selectively recognizes active arrestin2 ^20^. The mixtures were analyzed by size exclusion chromatography (SEC) (Figure 2D). Both CCR5 phosphopeptide 6P (red) or V2Rpp (green) induced a shift of the elution volume to ~2.3 ml from the ~2.6 ml observed for the inactive apo form (black), which indicates the formation of stable phosphopeptide•arrestin2•Fab30 complexes. An identical result was obtained for the peptide 3P2 (blue), which is only phosphorylated at the central T340, S342, and T343 residues, despite its more than four-fold lower affinity. In contrast, all other CCR5 phosphopeptides with similar low affinities as 3P2 resulted in mixtures between active and inactive arrestin populations. These results suggest that the 3P2 phosphorylation pattern pXpp is specific, necessary, and sufficient for inducing the active arrestin2 conformation as assayed by Fab30 binding, while the other 6P phosphosites only contribute to the overall affinity.

### Crystal structures of human arrestin2 in apo state and in complex with distinct CCR5 phosphopeptides

To obtain insights into structural changes of arrestin2 induced by phosphopeptide binding, two complexes of human arrestin2 (arrestin2^1-359^, lacking the C-terminal strand β20, see below) with the CCR5 4P and 6P phosphopeptides and the stabilizing Fab30 were prepared and their crystal structures determined at resolutions of 3.2 and 3.5 Å, respectively (Figure 3A, Table S1). Furthermore, also the crystal structure of full-length human arrestin2 was determined in its apo state at a resolution of 2.3 Å (Figure 3A). The latter is highly similar to the structure of bovine arrestin2 (PDB 1G4M) with the conserved fold composed of the two N- and C-terminal β-sandwich domains, the characteristic parallel-sheet between the N- and C-terminal strands 1/20, as well as a disordered C-terminal tail beyond residue 396 ^37^.

**Figure 3.**
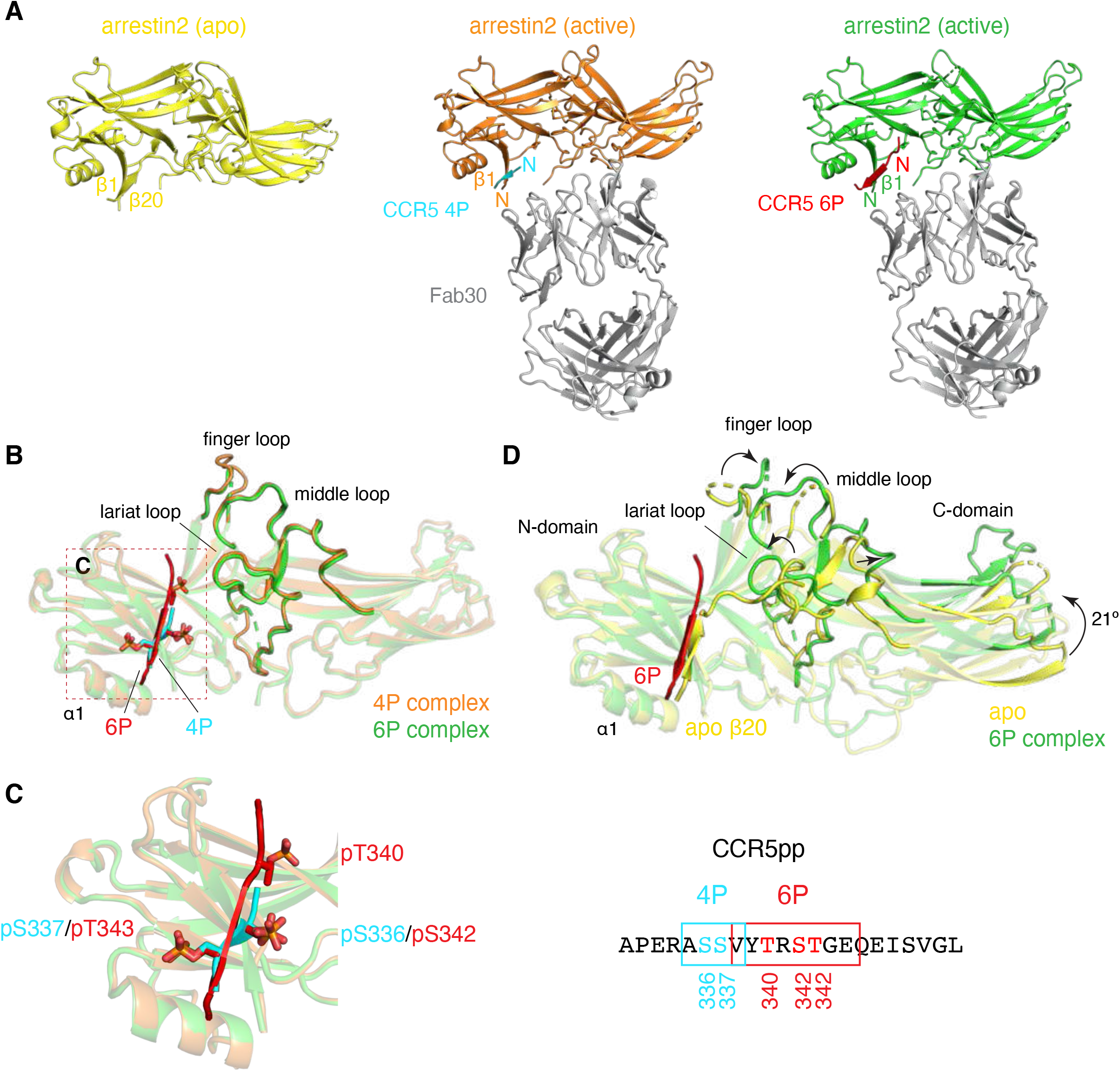
X-ray crystallographic structures of arrestin2 in apo form and in complexes with two CCR5 phosphopeptides. (A) Solved structures of apo arrestin2 (yellow), 4P•arrestin2 (arrestin2: orange, 4P: cyan), and 6P•arrestin2 (arrestin2: green, 6P: red) complexes. Both arrestin2 phosphopeptide complexes were stabilized with Fab30 (gray) (B) Overlay of the two arrestin2 complexes with the 4P (cyan) and 6P (red) phosphopeptides. (C) Detailed view of the peptide binding interfaces of both complexes (left) and alignment of the respective CCR5 C-terminal residues (right) showing a stable interaction in the electron density of the arrestin2 complexes. (D) Structural overlay of inactive, apo arrestin2 (yellow) and active arrestin2 (green) in complex with 6P (red). Salient arrestin2 conformational changes upon activation are indicated by black arrows.

While only parts of the phosphopeptides have well-defined electron densities in both phosphopeptide complex structures (Figure S3A), the phosphopeptide electron density is better defined in the 6P complex structure, indicating lower structural disorder in this region. For 6P, eight peptide residues (V338–E345) including the three phosphosites (pT340, pS342, pT343) have clearly defined density. In contrast, the electron density for 4P is reasonably defined only for four residues (A335–V338) including only two phosphosites (pS336, pS337) (Figure 3B). Both 6P and 4P coordinate as extended β-strands with the arrestin2 N-terminal strand β1 in an antiparallel manner with their observable phosphate groups interacting with the same arrestin2 residues (see below). Strikingly, however, there is a register shift (Figure 3C) such that the two consecutive phosphosites pS336, pS337 of the 4P complex take the positions of the two consecutive phosphosites pS342, pT343 in the 6P complex.

Both phosphopeptide complexes exhibit the hallmarks of arrestin2 activation, i.e., the formation of the anti-parallel intermolecular β-sheet between the arrestin strand β1 and the phosphopeptide, which replaces strand β20 of the intramolecular parallel β-sheet in the inactive apo arrestin2 structure, a twist of the C-domain relative to the N-domain of approximately 21°, and significant changes in the lariat, finger and middle loops (Figure 3D). The two phosphopeptides induced very minor differences in the orientations of Fab30 relative to arrestin2 (Figure S3B) and consequently also very minor differences in unit cell dimensions and crystal packing of their crystallized complexes despite identical space groups (Table S1). As compared to the 4P complex, the electron density of the 6P complex is less well-defined for arrestin2 residues 64-70 (center of finger loop) and 308-313 (at the end of the lariat loop). This may be caused by the variations in the crystal packing, but also indicates structural plasticity of these loops, which participate in receptor (finger loop) and clathrin (lariat loop) binding ^38^.

### Key CCR5 phosphorylation sites responsible for arrestin2 activation

The polar core of apo arrestin harbors a network of highly conserved ionic interactions formed by D26 and R169 in the N-domain, D290 and D297 on the lariat loop of the C-domain and R393 located in an extended stretch after the C-terminal strand β20 (Figure 4A). Together with a salt bridge between R25 and E389 and the hydrogen bonds forming the parallel β-sheet between strands β1 and β20, these strong interactions connect the two arrestin domains and stabilize the inactive conformation.

**Figure 4.**
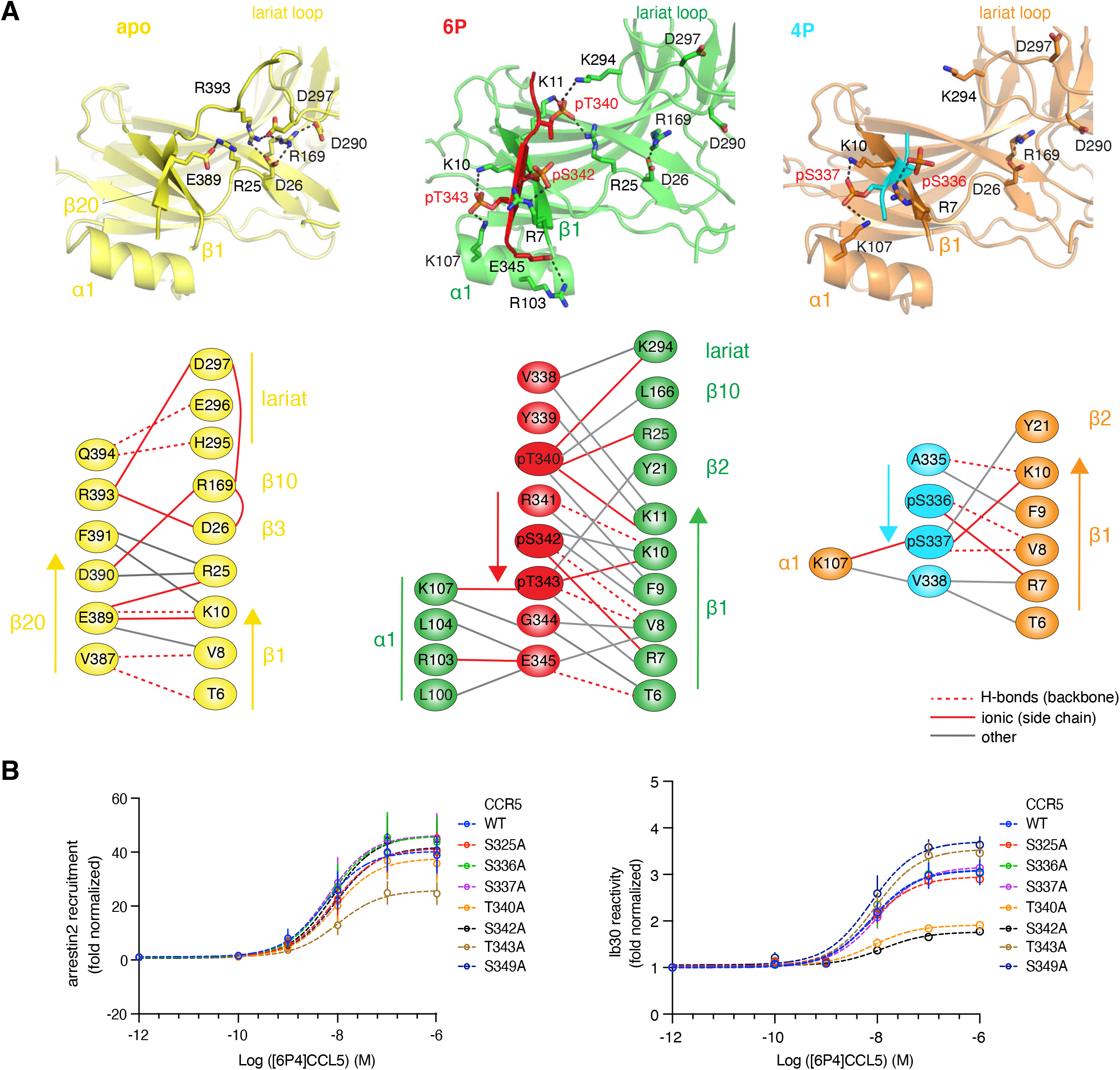
Detailed view of arrestin2 receptor C-terminal tail interactions. (A) Structural details of the arrestin2 N-domain in apo form (left) and in complexes with the CCR5 6P (center) and 4P phosphopeptides (right). Important residues stabilizing the inactive and active conformations are depicted in stick representation with key interactions as dashed lines. Upon arrestin2 activation by the peptides, the polar core is perturbed. Schematic diagrams of the key residue interactions are shown below each structural panel. The color coding follows Figure 3. (B) Super-agonist [6P4]CCL5-induced arrestin2 activation by CCR5 monitored with NanoBiT assays in HEK293 cells. Left: [6P4]CCL5-induced arrestin2 recruitment by wild-type CCR5 and seven S/A or T/A CCR5 C-terminal point mutants. Right: [6P4]CCL5-induced arrestin2 conformational changes monitored by the Ib30 assay for the same CCR5 constructs.

The central element of arrestin activation is the disruption of this polar core by the β-strand exchange of the arrestin strand β20 with the receptor phosphopeptide and the subsequent relocation of the gate loop ^20^. Arrestin2^1-359^, which was used for solving the crystal structures of the active complexes with the CCR5 phosphopeptides, lacks strand β20, thereby facilitating the formation of the intermolecular anti-parallel β-sheet to arrestin2 strand β1. In the 6P complex, the intermolecular β-sheet comprises CCR5 residues 341 to 344 (Figure 4A). The first visible phosphoresidue pT340 forms intermolecular salt bridges to R25, K11, and K294 (lariat loop), thereby apparently disrupting the salt bridges between R169 and D290/D297, pulling the lariat loop towards the N-domain, and forcing a twist of the C-domain. The 6P complex is further stabilized by extensive electrostatic interactions of CCR5 residues pS342, pT343, and E345 with arrestin2 residues R7, K10 on strand β1 and R103, K107 on helix α1. In addition, the phosphate of pS342 forms a salt bridge to R67 on Fab30 (Figure S3B).

While the 4P peptide also establishes an intermolecular anti-parallel β-sheet with strand β1 of arrestin2, there are fewer stable intermolecular contacts with no engagement of the arrestin2 lariat loop (Figure 4A). The two visible phosphoresidues, pS336 and pS337, form intermolecular salt bridges with arrestin2 residues R7, K10, and K107 in a similar way as the 6P phosphoresidues pS342 and pT343.

#### Cellular assays

The effect of individual CCR5 phosphorylation sites on arrestin2 recruitment and conformation was tested in a cellular context using several previously developed luciferase (NanoBiT) complementation assays ^39^ on seven S/A and T/A C-terminal point mutations of full-length CCR5.

For testing the direct recruitment of arrestin2 by CCR5 in response to [6P4]CCL5 stimulation, CCR5 fused to a small fragment (SmBiT) and arrestin2 fused to a large fragment (LgBiT) of luciferase (NanoBiT) were co-expressed in HEK293 cells and the complementation-induced luminescence was measured upon stimulation with the super-agonist [6P4]CCL5 (Figure 4B, S4A). With the exception of the strongly attenuated T343A, all other CCR5 mutants recruited arrestin2 with similar efficiency as wild-type CCR5. We attribute the reduced recruitment by T343A to the abolishment of the direct ionic interaction of the pT343 phosphate group with the side chain of the arrestin N-terminal residue R7 (Figure 4A), which also leads to a three-fold reduction in the phosphopeptide affinity as assayed by NMR (see above, Figure 2). The intrabody30 (Ib30, a single chain derivative of Fab30) specifically recognizes the activated conformation of arrestin2 ^40^. For testing the formation of the activated conformation, Ib30 fused to LgBiT and arrestin2 fused to SmBiT, and the respective CCR5 mutant were co-expressed and the cells stimulated by [6P4]CCL5 (Figure 4B, S4A). Whereas most mutations had only a moderate effect, the CCR5-T340A and S342A mutants almost completely abolished arrestin recognition by Ib30. For S342A this is expected, since pS342 directly contacts Fab30 in the 6P•arrestin2•Fab30 complex (Figure 3A). In contrast, the strong effect of T340A mutation apparently indicates a genuine change of the arrestin2 conformation, which strongly impedes recognition by Ib30. This agrees with the notion that pT340 is directly involved in the arrestin2 activating motion by pulling the lariat loop towards the N-domain (see above and below).

Finally, we also assayed endocytosis for the three CCR5 mutants (T340A, S342A and T343A) by co-expression with arrestin2 fused to SmBiT and the endofin FYVE domain, which targets early endosomes, fused to LgBiT (Figure S4C). In agreement with the arrestin2 recruitment assays, only CCR5-T343A significantly decreased endocytosis levels.

### The phosphorylation motif pXpp is responsible for stable arrestin recruitment

#### CCR5 vs V2R

The phosphorylation patterns of the V2R and its interactions with arrestin2 have been studied extensively by a range of biophysical and computational methods ^9,19,20,41,42^. Little is known about the interactions of other receptors with arrestin2. The present structures of the CCR5 C-terminal peptides in complex with arrestin2 provide an opportunity for a detailed comparison.

Both CCR5 6P•arrestin2•Fab30 and V2Rpp•arrestin2•Fab30 (PDB: 4JQI) complexes have an overall very similar organization (Figure S5). Despite V2Rpp containing five more phosphorylation sites than CCR5 6P, both peptides form a completely analogous antiparallel β-sheet between their central residues and strand β1 of arrestin2 (Figure 5A). This β-sheet is stabilized by an identical set of contacts between positively charged residues of arrestin2 and a central pXpp motif on the phosphopeptides (6P: pT340, pS342, pT343; V2Rpp: pT360, pS362, pS363). Similar to pT360 of V2Rpp, pT340 of CCR5 6P is oriented towards the connector region between arrestin’s N- and C-domains and forms salt bridges to K11 and R25 in the N-domain as well as K294 on the lariat loop. The latter interaction may constitute a main component of the force that reorients the arrestin domains upon activation. This notion is corroborated by Ib30-based functional studies, which show that both pT340 in CCR5 (Figure 4B) and pT360 in V2R ^41^ strongly impact the conformation of arrestin2. Thus, this first phosphoresidue of the pXpp motif appears crucial to drive arrestin activation. Of note, the interaction of this first phosphoresidue is not present in the 4P complex where only the consecutive pS336 and pS337 take the role of the last two phosphoresidues in the pXpp motif (Figure 4B). However, the presence of Fab30 and crystal packing may stabilize the active arrestin domain orientation even for this incomplete pXpp motif.

**Figure 5.**
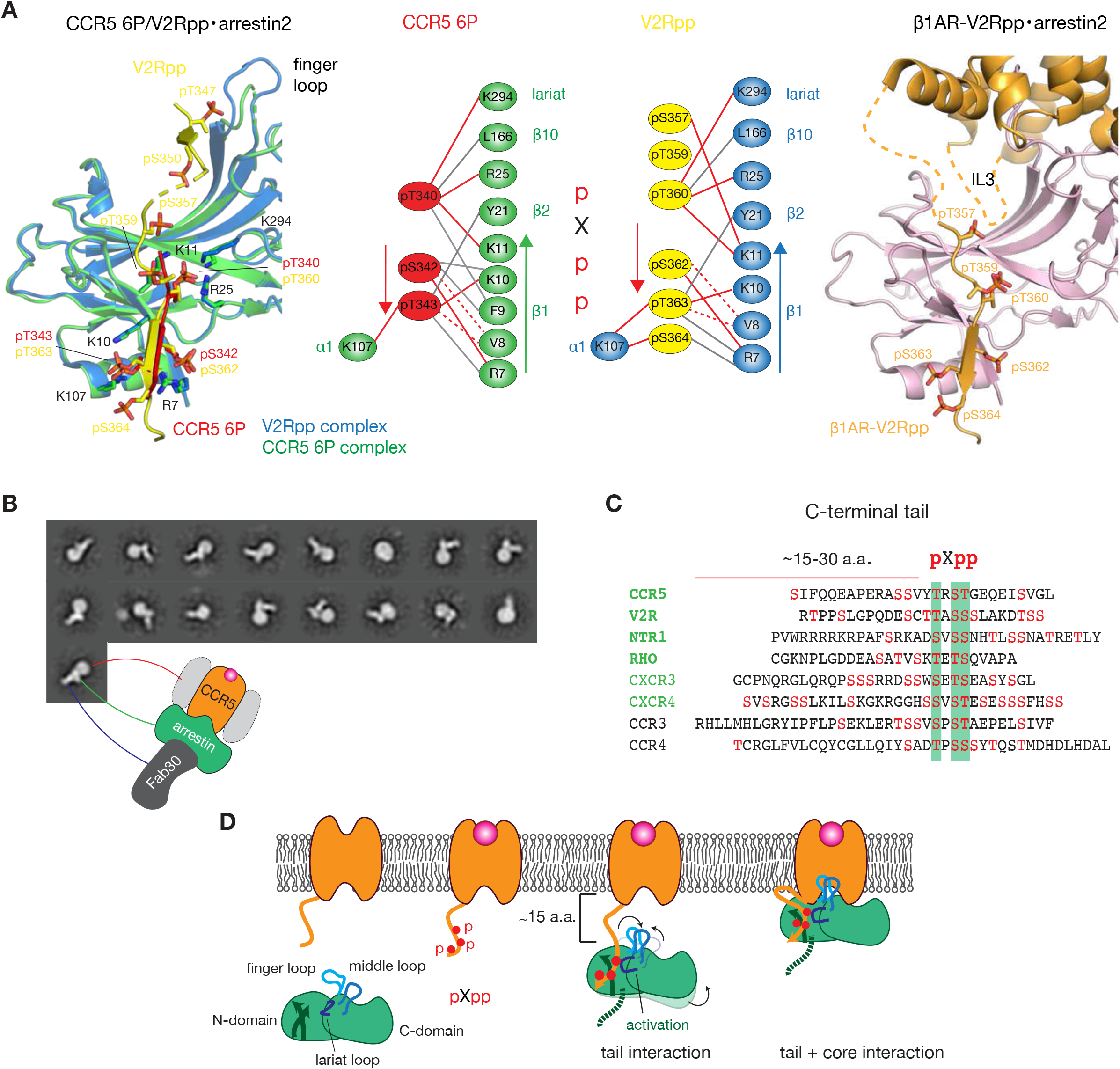
The pXpp motif is responsible for stable arrestin2 recruitment. (A) Left: Overlay of binding interfaces of arrestin2 with CCR5 6P (6P: red, arrestin2: green) and V2Rpp (V2Rpp: yellow, arrestin2: blue, PDB 4JQI). Phosphoresidues are shown in stick representation. Center: schematic diagrams of key 6P/V2R•arrestin2 interactions. Solid red lines represent polar interactions, dashed red lines H-bonds, gray lines other interactions. Right: V2Rpp•arrestin2 binding interface in the β1AR-V2Rpp•arrestin2 cryo-EM structure (PDB 6TKO). Unresolved parts of the ICL3 and C-terminal tail are depicted as dashed lines. (B) Single-particle analysis of negative-stain EM images of the [6P4]CCL5•CCR5•arrestin2•Fab30 complex. (C) Sequence alignment of C-terminal tails of several GPCRs (green), which are known to form a stable complex with arrestin2. Bold green indicates solved complex structures, light green functionally well-characterized receptors. Receptors indicated in black are examples of other not well characterized chemokine receptors harboring a pXpp motif. The GPCR C-terminal tail sequences were downloaded from the GPCRdb ^43^. The common pXpp motif (highlighted in green) is typically located about 15–30 residues downstream from receptor helix 8. (D) Schematic mechanism of arrestin2 (green) activation upon binding to an agonist(magenta)-stimulated class B GPCR (orange) containing a pXpp C-terminal cluster (red dots).

Besides these main interactions, further phosphosite interactions are observable in the V2Rpp complex, which are not present in the CCR5 6P complex: pT359 is involved in crystal contacts and pS357 as well as pS364 interact with the same arrestin residues as their respective adjacent pT360 and pT363. Remarkably, the N-terminal V2Rpp residues pT347 and pS350 form contacts with observable electron density to the arrestin2 finger loop and nearby residues (Figure 5A). However, these two phosphoresidues do not form visible interactions with arrestin2 in full-length receptor complexes such as the M2R-V2Rpp chimera•arrestin2 (PDB: 6U1N), the β1AR-V2Rpp chimera•arrestin2 (PDB: 6TKO) (Figure 5A) and the recently solved V2R•arrestin2 structures ^16^. Rather in these complexes, only the phosphosites beyond V2R residue 356 coordinate with the arrestin2 N-domain in an identical manner as in the V2Rpp and CCR5 6P complex structures. This is presumably due to steric constraints imposed by the direct binding of the receptor core to the arrestin2 finger loop ^14–16^. An inspection of the β1AR-V2Rpp chimera•arrestin2 structure (Figure 5A) shows that a flexible linker of about 15 residues connects the receptor helix 8 and the phosphorylation motif recognized by the arrestin2 N-domain.

To prove that wild-type CCR5 is able to engage arrestin2 robustly, we reconstituted a complex between arrestin2^1-393^, [6P4]CCL5, and GRK2-phosphorylated CCR5, for which the phosphorylation of each individual phosphosite had been verified by western blot analysis (see Methods). The complex was assembled on FLAG beads and stabilized by Fab30, washed extensively to remove excess arrestin and Fab30, and then further purified with SEC. Negative-stain EM of the complex and subsequent 2D classification revealed CCR5•arrestin2•Fab30 particles with arrestin in core-and tail-engaged as well tail-only-engaged arrangements. The latter may be caused by the dynamics of binding and/or heterogeneous phosphorylation of CCR5 (Figure 5B). These results prove that similar to NTR1 and V2R also GRK-phosphorylated CCR5 forms a stable complex with arrestin2.

## Discussion

### Generalization to other receptors

While our NMR data indicate that cumulative phosphorylation of the CCR5 C-terminus is important for arrestin2 affinity, the SEC data and Ib30 cellular assay show that a specific arrangement of phosphoresidues in a pXpp motif is needed for robust activation of arrestin2 and stable complex formation (Figures 2, 4). A comparison of the C-terminal sequences of structurally and functionally characterized GPCRs in the context of arrestin2 recruitment (Table S2) reveals that this pXpp motif is commonly found at a distance of 15 to 30 residues downstream of helix 8 (defined by the GPCRdb numbering scheme ^43^) (Figure 5C). This is the case for V2R, NTR1, and the chemokine receptors CXCR3 and CXCR4 (Figure 5C). A similar pattern occurs in many other chemokine receptors, such as CCR3 and CCR4 (Figures 5C, 6B). In agreement with these observations, the indicated phosphorylation sites have been suggested as key arrestin binding motifs in V2R ^41^, rhodopsin ^22^, and NTR1 ^18^. In summary (Figure 5D), this suggests that the pXpp cluster which follows a flexible linker of at least 15 amino acids after helix 8 is crucial for full arrestin2 engagement and stable arrestin2 complex formation. As judged from the high similarity of the solved CCR5 6P and V2Rpp complexes and the functional data, the first phosphoresidue of the pXpp cluster engages the lariat loop thereby triggering arrestin activation, whereas the last residue seems more important for overall arrestin2 recruitment (Figure 4B).

### Structural insights into arrestin isoform recognition

The arrestin2 and arrestin3 isoforms of non-visual arrestins have highly conserved sequences and similar three-dimensional structures. Both arrestins can desensitize GPCRs, however, their localization in cells differs to some extent. Whereas arrestin3 is localized in the cytoplasm, arrestin2 is found in both the cytoplasm and nucleus ^44,45^. Despite having similar interactions with client proteins, in many cases they play distinct roles in downstream outcomes ^8^.

Based on the characteristics of their agonist-dependent arrestin interaction, GPCRs have been separated into ‘class A’ and ‘class B’ subcategories ^7,46^. Class A receptors such as adrenergic, muscarinic, dopamine, μ-opioid, and 5-hydroxytryptamine receptors bind arrestin3 with higher affinity than arrestin2, but the respective complexes are transient and dissociate at or near the plasma membrane (Table S2). In contrast, class B receptors such as the angiotensin II type 1 receptor (AT1aR), NTR1, V2R rhodopsin, the complement C5a receptor, CCR5 and several others (Table S2) bind both arrestins with approximately equal, but higher affinity than class A receptors, and form long-lived arrestin complexes that traffic into endosomes.

#### Arrestin2 vs arrestin3 phosphopeptide complexes

To obtain insights into the specific recognition of both arrestin isoforms by GPCRs, we compared the structure of the arrestin2•CCR5 6P complex with that of arrestin3 in complex with the C-terminal ACKR3 phosphopeptide [ACKR3pp, PDB 6K3F ^21^, Figure 6A]. Similar to the CCR5 6P peptide, ACKR3pp binds in an extended conformation to the groove formed by the N-terminal arrestin β-sheet. However, its position is shifted towards the arrestin finger loop and it does not form an antiparallel β-sheet with the arrestin3 N-terminal strand β1. The arrestin3 binding groove also has a higher positive charge density than arrestin2, which extends towards the finger loop thereby explaining the engagement of ACKR3pp residue pS335 with this region. Both peptide positions overlap at the sites of pT340 (6P) and pT342 (ACKR3pp), interacting with a similar set of residues in arrestin2 and arrestin3, respectively. Interestingly, pS337 of CCR5, although not resolved in our structure and manually placed for comparison in Figure 6A would fit well into the arrestin3 binding interface and would form the same set of charge interactions as pT338 of ACKR3pp. In agreement with these observations, functional data on CCR5 indicate that an S337A mutant reduces both arrestin3 and arrestin2 binding to a similar extent, whereas arrestin3 binding is less reduced than arrestin2 binding for S342A and S349A mutants ^30^. This hints at a weaker role of CCR5 phosphoresidues beyond S337 in arrestin3 recruitment.

**Figure 6.**
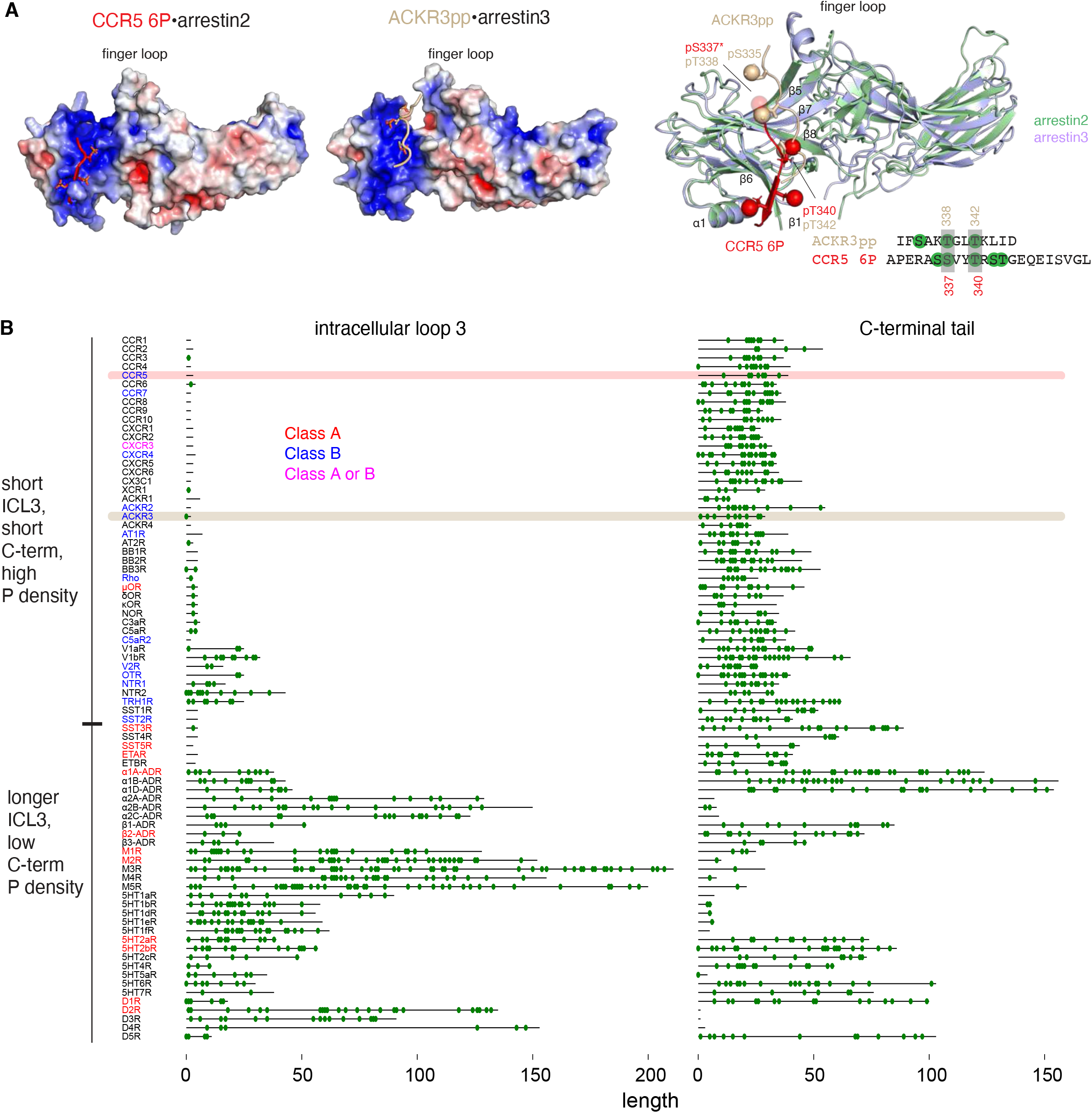
Structural differences in phosphopeptide recognition by arrestin2 and arrestin3. (A) Structural comparison of arrestin2 bound to the CCR5 6P phosphopeptide and arrestin3 bound to the ACKR3pp (PDB 6K3F). Left and center: arrestin electric surface charge densities (red: positive, blue: negative) of the 6P•arrestin2 and ACKR3pp•arrestin3 complexes. Right: superposition of the two complex structures (arrestin2: pale green, CCR5 6P: red, arrestin3: blue, ACKR3pp: wheat). Phosphates are shown as spheres (the position of the CCR5 pS337* phosphate is modeled). A sequence alignment is shown at the bottom with the overlapping phosphoresidue recognition sites indicated by gray boxes. (B) Sequence alignments of intracellular loops 3 (ICL3) and C-terminal tails of representative GPCRs. The amino acid sequences of GPCR ICL3 and C-terminal tails were downloaded from the GPCRdb ^43^. Potential phosphorylation sites (S and T) are shown as green circles. Receptors, which have been characterized for the type of arrestin binding (Table S2) are colored in red for class A, blue for class B, and purple for receptors, which exhibit isoform-dependent class A or B behavior. Sequences for CCR5 and ACKR3, for which the structures of arrestin complexes are shown in panel A, are highlighted.

#### Receptor ICL3 and C-terminal tail sequence analysis

Besides these structurally well-detected interactions between the GPCR C-terminal tails and arrestins, also intracellular loops (ILs) interact with arrestins ^12,14,18^, albeit they mostly have not been resolved. In particular, ICL3 is in very close proximity to arrestin (see e.g., β1AR-V2Rpp•arrestin2 complex, Figure 5A). To relate these structural findings to arrestin specificity we aligned the ICL3 and C-terminal tail sequences of GPCRs that are well-characterized with respect to arrestin interactions (Table S2) together with receptors from similar families (Figure 6B).

A visual inspection of the location of potential phosphorylation sites (S/T) and lengths of the IL3s and the C-terminal tails clearly puts the analyzed receptors into two distinct groups. The first group (Figure 6B, top) comprises receptors with very short IL3s (< ~5aa) containing few possible phosphorylation sites and short (~20–50 aa) C-terminal tails with dense clusters of serine and threonine residues often harboring the pXpp motif. Receptors of this group are almost all peptide-binding GPCRs including chemokine receptors with a number of them known to form stable arrestin complexes and characterized as class B receptors (Table S2). As indicated, both CCR5 and V2R belong to this group. The high density of potential phosphorylation sites and the specific recognition of the pXpp motif in this class of receptors may explain the higher affinity for arrestins and the long lifetime of the arrestin complexes.

The second group of receptors (Figure 6B, bottom) have longer to extremely long (>100 aa) IL3s and C-terminal tails with diverse lengths ranging from very short (< 10 aa, muscarinic, dopamine, and some 5-hydroxytryptamine receptors) to very long (~150 aa, α-adrenergic receptors). The density of potential phosphorylation sites within ICL3 and the C-terminal tail is lower for this second group than for the first group. The receptors in this second group, which have been characterized for arrestin binding (Table S2), interact with arrestins in a transient manner and in part have a preference towards arrestin3 as would be expected for class A receptors. Interestingly, the pXpp motif can also be found in the long ICL3 of some of these receptors such as M2R, which may enable specific arrestin2 binding also via ICL3 as evident from the Ib30 recognition of such complexes ^39^.

This diversity of C-terminal tail lengths and the low density of phosphorylation sites in the class A receptor group suggest that arrestin3 recruitment is less dependent on the exact position of phosphoresidues within the C-terminal tail than arrestin2 recruitment. The more extended positive surface of the binding groove may accommodate this diversity of phosphosites and lead to lower affinity/transient arrestin binding. Moreover, the structure of active arrestin3 in complex with ACKR3pp (Figure 6A) shows that the peptide interaction surface is very close to the finger and middle loops, which are engaged with the receptor core. Considering this close proximity of the arrestin3 binding interface to the receptor core, it is likely that both the C-terminal tail and ICL3 participate in the arrestin3 complex formation. Of note, D2R, which has a long ICL3, but no C-terminal tail after helix 8, recruits arrestin3 even in absence of GRK phosphorylation suggesting that arrestin3 recruitment might be less dependent on phosphorylation ^47^.

## Conclusion

In conclusion our structural and biophysical analysis of the interactions between the CCR5 phosphorylated C-terminus and arrestin2 has identified a key pXpp motif for arrestin2 recruitment and activation, which appears to be conserved across many class B GPCRs such as V2R, NTR1, rhodopsin, CXCR3 and CXCR4. An analysis of GPCR ICL3 and C-terminal phosphorylation sites and their class A or B arrestin interaction behavior together with a structural comparison of arrestin2 and arrestin3 phosphopeptide binding modes revealed salient sequence features for arrestin2 and arrestin3 isoform specificity. This may provide a framework for a more rigorous characterization of other GPCR arrestin systems, which are needed to obtain a comprehensive picture of the diverse arrestin binding modes and their relation to GPCR phosphorylation.

## Methods

### Peptide synthesis

Phosphorylated peptides corresponding to the last 22 residues of human CCR5 receptor (3P1, 3P2, 4P, 5P1, 5P2, 6P) or the last 29 residues of the V2 receptor (V2Rpp) were obtained from the Tufts University Core Facility for peptide synthesis. The non-phosphorylated CCR5 peptide (0P) was obtained from GenScript and contained a biotinylated N-terminus.

### Constructs

The genes encoding wild-type, full-length human CCR5 with a C-terminal 3C cleavage site followed by a FLAG tag for expression in the baculovirus Sf9 insect cell system and [6P4]CCL5 for expression in *E. coli* have been described before ^23,48^.

The plasmid encoding GPCR kinase subtype 2 used for the receptor phosphorylation in insect cells (GRK2-CAAX) was a gift from Robert Lefkowitz (Addgene plasmid #166224 ^49^). Full-length arrestin2^1-418^ (C150L, C242V, C251V, C269S) in vector pET-28a (+) was obtained from GenScript. The complete construct contained an N-terminal hexahistidine tag followed by a TEV cleavage site (ENLYFQG) and the arrestin2 sequence (bold):

MGSSHHHHHHSSGENLYFQG**MGDKGTRVFKKASPNGKLTVYLGKRDFVDHIDLVDPVDGVVLVDPEYLKERRVYVT LTCAFRYGREDLDVLGLTFRKDLFVANVQSFPPAPEDKKPLTRLQERLIKKLGEHAYPFTFEIPPNLPCSVTLQPGPEDTGKACGVDYEVKAFLAENLEEKIHKRNSVRLVIRKVQYAPERPGPQPTAETTRQFLMSDKPLHLEASLDKEIYYHGEPISVNVHVTNNTNKTVKKIKISVRQYADIVLFNTAQYKVPVAMEEADDTVAPSSTFSKVYTLTPFLANNREKRGLALDGKLKHEDTNLASSTLLREGANREILGIIVSYKVKVKLVVSRGGLLGDLASSDVAVELPFTLMHPKPKEEPPHREVPENETPVDTNLIELDTNDDDIVFEDFARQRLKGMKDDKEEEEDGTGSPQLNNR**

The truncated arrestin2^1-359^ and arrestin2^1-393^ constructs were obtained by introducing stop codons (TAA) into arrestin2^1-418^ at position 360 and 394, respectively, via standard QuickChange polymerase chain reactions. For crystallization of the apo form, arrestin2^1-418^ was obtained from a DNA construct cloned into a pGEX vector harboring an N-terminal GST tag followed by an HRV-3C cleavage site.

### Protein expression and purification

#### CCR5

Non-phosphorylated wild-type full-length CCR5 and super-agonist chemokine [6P4]CCL5 were expressed in Sf9 cells and *E. coli*, respectively, and purified according to previous protocols ^23,48^.

Phosphorylated CCR5 was obtained by co-expression with untagged GRK2-CAAX in Sf9 insect cells following a protocol described for the phosphorylation of the β2-adrenergic receptor ^6^ and optimized as follows. Once the viability dropped to ~85–90% [44 hour post-infection (hpi)], cells were stimulated by addition of 500 nM [6P4]CCL5. After 2 h incubation at 37 °C, cells were harvested and kept at −80 °C until further use. The phosphorylation level was assayed using western blot analysis with phospho-specific CCR5 antibodies. Membrane preparation and receptor purification were carried out as previously described ^48^.

#### Arrestin

Arrestin2^1-418^, arrestin2^1-393^ and arrestin2^1-359^ were expressed in *E. coli* BL21 (DE3) strain cultured in Lysogeny broth (LB). For the preparation of the deuterated ^15^N-labeled arrestin2^1-393^ NMR sample, cells were grown in D_2_O/^15^NH_4_Cl M9 minimal medium. Cells were grown at 37 °C until the optical density at 600 nm reached 0.7–0.8. Thereafter protein expression was induced by the addition of 25 μM isopropyl-^D^-thiogalactopyranoside and the temperature lowered to 18 °C for an overnight expression, after which the cells were harvested by centrifugation. Proteins were purified on a Ni-NTA HiTrap HP column (GE Life Sciences) and the His tag was removed by overnight cleavage with TEV protease (homemade). The cleaved protein was further separated from impurities by a reverse IMAC step on a Ni-NTA HiTrap HP column, followed by concentration in a Vivaspin 20 concentrator [10-kDa MWCO (molecular weight cutoff)] and final gel filtration step on a HiLoad 16/600 Superdex 200 pg gel filtration column (GE Healthcare) equilibrated with 20 mM HEPES, 150 mM NaCl, pH 7.4 (SEC buffer I). The protein purity was confirmed by SDS-PAGE.

The GST-tagged arrestin2^1-418^ was expressed under the same conditions, purified using a GST HiTrap column, followed by removal of the GST tag by homemade PreScission protease and a final gel filtration step on a HiLoad 16/600 Superdex 200 pg gel filtration column.

### Preparation of Fab30

Fab30 was purified as described previously ^20^, with slight modifications. Briefly, overnight a primary culture of Fab30-transformed *E. coli* M55244 strain was inoculated into 1 L 2x YT media (Himedia, Cat. no. G034) and allowed to grow for 8 hours at 30 °C. After 8 hours the culture was pelleted down and redissolved in 1 L CRAP media [7 mM (NH4)2SO4, 14 mM KCl, 2.4 mM sodium citrate, 5.4 g/L yeast extract, 5.4 g/L casein hydrolyzates, 0.11 M MOPS buffer pH 7.3, 0.55% (w/v) glucose, 7 mM MgSO_4_] grown for 16-18 h at 30 °C. Harvested cells were lysed in 20 mM HEPES (SRL, Cat. no. 63732), 100 mM NaCl, 0.5 mM MgSO_4_, 0.5% Triton-X 100, pH 8.0. Cell debris was separated by high-speed centrifugation. After loading the cell lysate onto a Protein L beads (Capto™ L, GE Healthcare, Cat. no. 17-5478-02) column, nonspecific proteins were removed through extensive washing [20 mM HEPES, 100 mM NaCl, pH 8.0]. Bound protein was eluted in 0.1 M acetic acid, pH 3.0, and neutralized with 1 M HEPES, pH 8.0. Eluted protein was desalted using a PD10 column (GE Healthcare, Cat. no. 17085101) in 20 mM HEPES, 150 mM NaCl, pH 8.0. The purified protein was stored at −80 °C in 10% glycerol until further use.

### Western blot analysis of CCR5 phosphorylation by phospho-specific CCR5 antibodies

All the following steps were performed at room temperature. 10 μL of non-phosphorylated or phosphorylated FLAG-purified CCR5 were separated by SDS-PAGE and transferred to a nitrocellulose membrane. Blots were then blocked with 1% BSA in Tris-Buffered Saline-Tween (TBST) for 1 h and then incubated with different primary (rabbit polyclonal) phospho-CCR5 antibodies [pS336/pS337-, pS342-, pT340-CCR5 (all 7TM antibodies) and pS349-CCR5 (Thermo Fisher)] at 1:2000 dilution for 1 h. Blots were washed twice for 5 min with TBST and incubated with HRP-coupled anti-rabbit secondary antibody (Thermo Fisher) at 1:5000 dilution for 1 h in the dark. Blots were then washed three times for 5 min with TBST and developed using western blotting substrate chemiluminescent detection.

### Mass spectrometry

#### Sample preparation

The samples of non-phosphorylated and phosphorylated FLAG-purified CCR5 were prepared as technical triplicates. For this, 5 μg of the receptor was reduced and alkylated for 10 min at 95 °C in 50 μl of 1% sodium deoxycholate, 0.1 M ammonium bicarbonate, 10 mM TCEP, 15 mM chloroacetamide, pH 8.3. The sample was split into two, and the two halves were digested with either Sequencing Grade Modified Trypsin or endoproteinase Glu-C (both Promega, Madison, Wisconsin, enzyme:receptor = 1:50 w/w) for 12 h at 37 °C. The samples were then acidified by the addition of 50 mM HCl (from 2 M HCl stock) and incubated for 15 min at 37 °C. Subsequently, the precipitated detergent was removed by centrifugation at 10,000g for 15 min and the peptides were separated from the reaction mixture using a C18 spin column (BioPureSPN MINI, The Nest Group, Inc.) according to the manufacturer’s instructions. Samples were then dried under vacuum and stored at −80 °C until further use.

#### LC-MS analysis

For LC-MS analysis, the dried peptide samples were solubilized at a concentration of 1 pmol/μl in 98% water, 2% acetonitrile, 0.15% formic acid. 5 μl of each sample was then subjected to LC-MS analysis by a Q Exactive Plus mass spectrometer fitted with an EASY-nLC 1000 liquid chromatography system (both Thermo Fisher Scientific). Peptides were resolved using an EasySpray RP-HPLC column (75 μm × 25 cm) at a flow rate of 0.2 μL/min and a pre-column setup under a linear gradient ranging from 5% buffer B (80% acetonitrile, 0.1% formic acid in water) in 95% buffer A (0.1% formic acid in water) to 45% buffer B over 60 minutes. The mass spectrometer was operated in DDA (data-dependent acquisition) mode with a total cycle time of approximately 1 s. Each MS1 scan was followed by high-collision-dissociation (HCD) of the 20 most abundant precursor ions with the dynamic exclusion set to 5 seconds. For MS1, 3e6 ions were accumulated in the Orbitrap over a maximum time of 25 ms and scanned at a resolution of 70,000 FWHM (at 200 m/z). MS2 scans were acquired at a target setting of 1e5 ions, maximum accumulation time of 110 ms and resolution of 17,500 FWHM (at 200 m/z). Singly charged ions, ions with charge state ≥ 6 and ions with unassigned charge state were excluded from triggering MS2 events. The normalized collision energy was set to 27%, the mass isolation window to 1.4 m/z, and one microscan was acquired for each spectrum.

In addition to the DDA LC-MS analysis, a targeted MS analysis was carried out focusing on the peptides with the phosphorylation sites of interest. For this, the sequence of CCR5_HUMAN was downloaded from uniprot.org (download 2022/10/22), imported into the Skyline software (https://skyline.ms/project/home/software/Skyline/begin.view), and the corresponding peptides manually phosphorylated at the expected sites of phosphorylation. The peptide ion masses containing 2^+^ and 3^+^ ions were exported as a mass isolation list from Skyline v21.2 and imported to the MS acquisition software. Targeted LC-MS analysis was carried out using the same settings as for DDA LC-MS with the following changes: the resolution of MS1 scans was reduced to 35,000 FWHM (at 200 m/z), the AGC target for MS2 scans was set to 3e6, the maximal fill time to 50 ms, and the mass isolation window to 0.4 m/z.

#### MS data analysis

The acquired raw files were converted to the mascot generic file (mgf) format using the msconvert tool [part of ProteoWizard, version 3.0.4624 (2013-6-3)]. Using the MASCOT algorithm (Matrix Science, Version 2.4.1), the mgf files were searched against a decoy database containing normal and reverse sequences of the predicted UniProt entries of *Spodoptera frugiperda* (www.ebi.ac.uk, release date 2020/10/22), the protein CCR5_HUMAN and commonly observed contaminants (in total 56,642 sequences) generated using the SequenceReverser tool from the MaxQuant software (version 1.0.13.13). The precursor ion tolerance was set to 10 ppm and fragment ion tolerance was set to 0.02 Da. The search criteria were set to requiring full trypsin specificity (cleavage after lysine or arginine unless followed by proline) and GluC specificity (cleavage after aspartate or glutamate unless followed by proline). At most 3 miscleavages were allowed, carbamidomethylation (C) was set as fixed modification, and phosphorylation (STY), oxidation (M) and acetylation (protein N-terminus) were set as variable modifications. The database search results were imported into Scaffold (version 5.1.0) and filtered to 1% FDR (false discovery rate) on the protein and peptide level using the built-in LFDR algorithm.

### Trypsin proteolysis assay

Arrestin2 trypsin proteolysis assays were carried out by first incubating 100 μL of 20 μM arrestin2^1-418^ with a 3-molar excess of the respective phosphopeptide at room temperature for 5-10 min in SEC buffer I. Thereafter, 1 ng of Trypsin Gold (Promega) was added, and the mixture incubated at 35 °C under gentle shaking (500 rpm). Samples for SDS-PAGE (10 μL sample mixed with 10 μL of 4x SDS loading buffer) were taken at 0, 5, 10, 20, 30, 60 and 90 min. The reaction was quenched by boiling the samples for 10 min at 95 °C. Samples were then loaded on 4-20% precast gradient gels and visualized with Instant Blue protein stain (Abcam). Control reactions were run with apo arrestin2^1-418^ and arrestin2^1-418^ incubated with the phosphopeptide 0P.

### NMR titrations

^15^N-, ^2^H-labeled arrestin2^1-393^ (50 μM) NMR samples were prepared in SEC buffer I supplemented with 5% D_2_O and 0.03% NaN_3_ as 270-μl volumes in Shigemi tubes. The phosphopeptides were titrated into these samples to concentrations of 0–500 μM and the interactions monitored by ^1^H-^15^N HSQC-TROSY spectra recorded on a Bruker AVANCE 14.1 T (600 MHz) spectrometer equipped with a TCI cryoprobe at 303 K.

NMR data were processed with NMRPipe ^50^ and analyzed with NMRFAM-SPARKY ^51^. K_D_ values were obtained by nonlinear least-squares fitting using Matlab (Matlab_R2021b, MathWorks, Inc.) and the following equation:

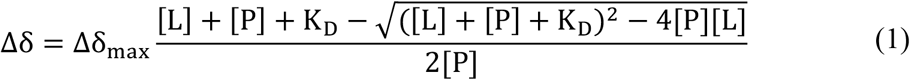

where Δδ = δ_apo_-δ_bound_ is the difference in the ^1^H^N^ or ^15^N arrestin2 chemical shift, δ_max_ is the difference between apo and the fully ligand-bound state, and [P] and [L] are the total protein (arrestin2) and ligand (phosphopeptide) concentrations, respectively.

### CCR5 phosphopeptide•arrestin2•Fab30 complex formation and purification

Arrestin2^1-359^ (30 μM) was incubated with a 5-molar excess of CCR5 phosphopeptides for 1 h on ice in SEC buffer II (20 mM HEPES, 150 mM NaCl, pH 6.9). Then, a 1.2-molar excess of Fab30 was added, followed by incubation for 1.5 h in the cold room. Samples were then concentrated in an Amicon concentrator (MWCO 50 kDa) and separated by SEC using a self-packed 4-ml S200 10/300 SEC column (length 25 mm, diameter 4.6 mm) and monitoring protein absorbance at 280 nm. The complex quality of each SEC fraction was evaluated by SDS-PAGE and visualized with Instant Blue protein stain. Fractions showing a fully formed complex were pooled.

### X-ray crystallography

Both apo arrestin2 and its phosphopeptide/Fab30 complexes were crystallized by sitting drop vapor diffusion at room temperature from 1:1 mixtures of protein in SEC buffer II and crystallization buffer. Respective protein concentrations before mixing and crystallization buffers were: (i) 10 mg/ml apo arrestin2^1-418^, 100 mM magnesium formate dihydrate, 100 mM bis-tris, 15% PEG3350, pH 7.0 and (ii) ~4-6 mg/ml arrestin2^1-359^•Fab30•CCR5 phosphopeptide, 100 mM magnesium formate dihydrate, 100 mM bis-tris, 15% PEG3350, pH 7.0. In both cases, crystals formed and reached their final size within 24–48 h. Thereafter, they were quickly soaked in the crystallization buffer mixed with 20–25% ethylene glycol, and then flash-frozen in liquid nitrogen.

Diffraction data were collected at the Swiss Light Source, Paul Scherrer Institute, Villigen, Switzerland at beamline X06DA and X06SA, processed with XDS ^52^ and scaled using Aimless (Evans 2006). For the 6P•arrestin2^1-359^•Fab30 complex, a single data set was collected. For the 4P•arrestin2^1-359^•Fab30 complex two data sets from the same crystal were merged, and for the apo arrestin2^1-418^ four data sets of three different crystals were integrated. All structures were determined by molecular replacement with PHASER (McCoy, Grosse-Kunstleve et al. 2007) contained in the CCP4 software package ^53^, using 4JQI ^20^ for complexes and 1G4M ^37^ for apo arrestin2 as search models. Models building was performed with COOT (Emsley and Cowtan 2004) and refinement with BUSTER-TNT ^54^ and PHENIX ^55^. The final models were evaluated with MolProbity (Chen, Arendall et al. 2010) and visualized with PyMOL (DeLano 2002). Data and refinement statistics are summarized in Table S1.

### [6P4]CCL5•CCR5•arrestin2 complex formation and negative-stain EM analysis

Membranes containing phosphorylated CCR5 from 1 L Sf9 cell culture were solubilized in 50 mM HEPES, 400 mM NaCl, 0.5% LMNG, pH 7.4 and incubated with 1 mL M2 anti-FLAG resin in the cold room overnight. To form the [6P4]CCL5•CCR5•arrestin2 complex, the resin was then incubated with ~3-5 μM arrestin2^1-393^ and 5 μM [6P4]CCL5 for 1 h. The complex was then stabilized by the addition of a 1.2-fold excess of Fab30, followed by a further 1-hour incubation. Thereafter, the resin was packed into a column and washed with 10 column volumes (CV) wash buffer (25 mM HEPES, 150 mM NaCl, 10 % glycerol, 0.01% LMNG, pH 7.4). Then the complex was eluted with 3 CV elution buffer (25 mM HEPES, 150 mM NaCl, 0.01% LMNG, 0.2 mg/ml FLAG peptide, pH 7.4) and further purified by SEC using a Superdex 200 Increase 10/300 GL column preequilibrated with SEC buffer III (10 mM HEPES, 150 mM NaCl, 0.01% LMNG, pH 7.4). The quality of each SEC fraction was assessed by SDS-PAGE. Fractions showing good complex integrity and purity were combined.

For negative-stain EM, the complex was diluted to 0.05 mg/ml in SEC buffer III. 5 μl of this solution was then applied onto freshly glow-discharged carbon-coated 300-mesh copper grids (produced in-house) and blotted with filter paper. The grids were stained with 2% (w/v) uranyl acetate for 30 s and imaged at a magnification of x135,000 on a FEI Tecnai G2 Spirit TEM operated at 80 kV and equipped with an EMSIS Veleta camera. 2D classification of single particles was carried out with CryoSPARC v.3.1.

### NanoBiT assays

The receptor constructs including wild-type CCR5 and all phosphosite mutants were synthesized from GenScript and subcloned in pcDNA3.1(+) vector with an N-terminal FLAG tag. For receptor-based NanoBiT assay, receptor constructs bearing a carboxyl-terminus SmBiT spaced with a flexible linker were cloned in the lab using enzymes *Kpn*I and *Sma*I in pCAGGS vector.

For assessing agonist-induced arrestin recruitment, trafficking, and conformational variability of CCR5^WT^ and phosphosite mutants, a luciferase enzyme-linked complementation-based assay (NanoBiT assay) was used following the protocol described earlier ^56^. For arrestin2^1-418^ recruitment, receptor-SmBiT (1.5 µg) and LgBiT-arrestin2^1-418^ (1.5 µg) constructs were used (Figure 4B) to transfect HEK293 cells using polyethylenimine (PEI) with DNA:PEI ratio as 1:3. Similarly, for arrestin2^1-418^ endosomal trafficking, receptor constructs (3 µg), SmBiT-arrestin2^1-418^ (3.5 µg) and LgBiT-FYVE (3.5 µg) were used (Figure S3). Assessing the conformational diversity of arrestin2^1-418^ bound to receptors was done by transfecting cells with receptors (3 µg), SmBiT-arrestin2^1-418^ (2 µg) and LgBiT-Ib30 (5 µg) (Figure 4B). After transfection, all the NanoBiT-based assays follow a common set of steps. Briefly, 16 h post-transfection cells were trypsinized and harvested, followed by resuspension in assay buffer containing 1x HBSS (Gibco, Cat. no. 14065-056), 0.01% bovine serum albumin (BSA, SRL, Cat. no. 83803), 5 mM 4-(2-hydroxyethyl)-1-piperazineethanesulfonic acid (HEPES), pH 7.4 with 10 μM coelenterazine (GoldBio, Cat. no. CZ05). Afterward, cells were seeded at a density of 1×10^5^ cells well^-1^ in a white 96-well plate and incubated for 1.5 h at 37 °C followed by 30 min at room temperature. After incubation baseline luminescence was measured using a multimode plate reader and then cells were stimulated with varying doses of [6P4]CCL5 followed by measurement of luminescence signal for 10-20 cycles and average data from 5^th^ to 10^th^ cycle were analyzed and presented using GraphPad Prism 9 (v3) software.

### Receptor surface expression assay

In order to measure the surface expression of the receptors in different assays, we used a previously described whole cell-based surface ELISA assay ^57^. Post 24 h of transfection, cells were seeded into a 0.01% poly-D-Lysine precoated 24-well plate at a density of 2×10^5^ cells per well and allowed to adhere and grow for 24 h. The next day, cells were washed with ice-cold TBS, fixed with 4% PFA (w/v in TBS) on ice for 20 min. After fixing, cells were washed again three times with TBS and incubated in 1% BSA prepared in TBS at room temperature for 1.5 h. Thereafter, the cells were incubated with anti-FLAG M2-HRP antibody (Sigma, Cat. no. A8592) (1:5000, 1 h at room temperature) followed by three washes with 1% BSA in TBS. The plates were developed with TMB-ELISA substrate (Thermo Fisher Scientific, Cat. no. 34028) until the light blue color appeared. The reaction was quenched by transferring 100 μl of the colored solution to another 96-well plate already having 100 μl of 1 M H_2_SO_4_, and the absorbance was recorded at 450 nm. For normalization of the ELISA reading with the total cell content of each well, cells were incubated with 0.2% (w/v) Janus Green (Sigma, Cat. no. 201677) for 15 min at room temperature after washing twice with TBS. For removing excess stain cells were washed thoroughly with water followed by developing the satin with the addition of 800 μl of 0.5 N HCl in each well, of which 200 μl of this solution was transferred to a 96-well plate for measuring the absorbance at 595 nm. The ELISA signal was normalized from the A450/A595 ratio and the values were plotted using the GraphPad Prism 9 (v3).

## Data availability

The following structural models for human arrestin2 have been deposited in the Protein Data Bank: 4P•arrestin2•Fab30 complex (PDB 8AS2), 6P•arrestin2•Fab30 complex (PDB 8AS3) and apo arrestin2 (PDB 8AS4). The CCR5 phosphoproteomics data have been deposited in the ProteomeXchange repository (PXD036220).

## Acknowledgments

This work was supported by the Swiss National Science Foundation (grants 201270 and IZLIZ3-200298 to S.G., 179323 to T.M.), the Indo-Swiss grant from the Department of Biotechnology to A.K.S (IC-12044(11)/4/2021-ICD-D), and by a Fellowship for Excellence by the Biozentrum Basel International PhD Program to I.P. We gratefully acknowledge Dr. A. Schmidt and U.K. Lanner (Biozentrum Proteomics Core facility) for recording and analyzing proteomics data on phosphorylated CCR5, C. Alampi and Dr. M. Chami (Biozentrum, BioEM facility) for their support with electron microscopy, the beamline staff at Swiss Light Source (PSI) for crystallographic data collection and support at beamlines X06SA and X06DA, Dr. H. Dwivedi-Agnihotri and Dr. M. Baidya (IIT) for their help with cellular assays, M. Rogowski and I. Hertel (Biozentrum) for expression and purification of [6P4]CCL5 and proteases.

## Author contributions

P.I., I.P., V.P., A.K.S., and S.G. conceived the study. I.P., P.I., A.R., V.P. and M.B. designed, expressed and purified proteins. P.I. prepared phosphorylated CCR5 samples and analyzed respective data. I.P. recorded NMR experiments and analyzed NMR data. R.P.J. and V.P. crystallized apo arrestin2. R.P.J. and P.I. crystallized arrestin2 in complex with 4P and 6P CCR5 phosphopeptides. P.I., I.P., S.G., and A.K.S. analyzed the structures with input from R.P.J. and T.M.. P.I. prepared the sample for negative stain EM, recorded and processed the EM data. A.R. designed the cellular experiments with guidance from A.K.S., P.S., and A.R. generated all the constructs and mutants for the cellular assays and performed their validation. P.I., I.P. and S.G. wrote the manuscript with input from A.K.S..

## Declaration of interests

The authors have no competing interests.

## Notes

### Competing Interest Statement

The authors have declared no competing interest.

## References

1. Lefkowitz, R.J., and Shenoy, S.K. (2005). Transduction of Receptor Signals by ß-Arrestins. Science 308, 512–517. 10.1126/science.1109237.

2. Gurevich, V.V., and Gurevich, E.V. (2019). GPCR Signaling Regulation: The Role of GRKs and Arrestins. Frontiers in Pharmacology 10. 10.3389/fphar.2019.00125.

3. Cahill, T.J., Thomsen, A.R.B., Tarrasch, J.T., Plouffe, B., Nguyen, A.H., Yang, F., Huang, L.-Y., Kahsai, A.W., Bassoni, D.L., Gavino, B.J., et al. (2017). Distinct conformations of GPCR–β-arrestin complexes mediate desensitization, signaling, and endocytosis. Proceedings of the National Academy of Sciences 114, 2562–2567. 10.1073/pnas.1701529114.

4. Kumari, P., Srivastava, A., Banerjee, R., Ghosh, E., Gupta, P., Ranjan, R., Chen, X., Gupta, B., Gupta, C., Jaiman, D., et al. (2016). Functional competence of a partially engaged GPCR–β-arrestin complex. Nat Commun 7, 1–16. 10.1038/ncomms13416.

5. Lohse, M.J., and Hoffmann, C. (2014). Arrestin Interactions with G Protein-Coupled Receptors. In Arrestins - Pharmacology and Therapeutic Potential Handbook of Experimental Pharmacology., V. V. Gurevich, ed. (Springer), pp. 15–56. 10.1007/978-3-642-41199-1_2.

6. Shukla, A.K., Westfield, G.H., Xiao, K., Reis, R.I., Huang, L.-Y., Tripathi-Shukla, P., Qian, J., Li, S., Blanc, A., Oleskie, A.N., et al. (2014). Visualization of arrestin recruitment by a G-protein-coupled receptor. Nature 512, 218–222. 10.1038/nature13430.

7. Smith, J.S., and Rajagopal, S. (2016). The β-Arrestins: Multifunctional Regulators of G Protein-coupled Receptors. J Biol Chem 291, 8969–8977. 10.1074/jbc.R115.713313.

8. Srivastava, A., Gupta, B., Gupta, C., and Shukla, A.K. (2015). Emerging Functional Divergence of β-Arrestin Isoforms in GPCR Function. Trends in Endocrinology & Metabolism 26, 628–642. 10.1016/j.tem.2015.09.001.

9. Latorraca, N.R., Masureel, M., Hollingsworth, S.A., Heydenreich, F.M., Suomivuori, C.-M., Brinton, C., Townshend, R.J.L., Bouvier, M., Kobilka, B.K., and Dror, R.O. (2020). How GPCR Phosphorylation Patterns Orchestrate Arrestin-Mediated Signaling. Cell 183, 1813–1825.e18. 10.1016/j.cell.2020.11.014.

10. Tobin, A.B. (2008). G-protein-coupled receptor phosphorylation: where, when and by whom. British Journal of Pharmacology 153, S167–S176. 10.1038/sj.bjp.0707662.

11. Nobles, K.N., Xiao, K., Ahn, S., Shukla, A.K., Lam, C.M., Rajagopal, S., Strachan, R.T., Huang, T.-Y., Bressler, E.A., Hara, M.R., et al. (2011). Distinct Phosphorylation Sites on the β2-Adrenergic Receptor Establish a Barcode That Encodes Differential Functions of β-Arrestin. Science Signaling. 10.1126/scisignal.2001707.

12. Zhou, X.E., He, Y., de Waal, P.W., Gao, X., Kang, Y., Van Eps, N., Yin, Y., Pal, K., Goswami, D., White, T.A., et al. (2017). Identification of Phosphorylation Codes for Arrestin Recruitment by G Protein-Coupled Receptors. Cell 170, 457–469.e13. 10.1016/j.cell.2017.07.002.

13. Cao, C., Barros-Álvarez, X., Zhang, S., Kim, K., Dämgen, M.A., Panova, O., Suomivuori, C.-M., Fay, J.F., Zhong, X., Krumm, B.E., et al. (2022). Signaling snapshots of a serotonin receptor activated by the prototypical psychedelic LSD. Neuron. 10.1016/j.neuron.2022.08.006.

14. Staus, D.P., Hu, H., Robertson, M.J., Kleinhenz, A.L.W., Wingler, L.M., Capel, W.D., Latorraca, N.R., Lefkowitz, R.J., and Skiniotis, G. (2020). Structure of the M2 muscarinic receptor–β-arrestin complex in a lipid nanodisc. Nature 579, 297–302. 10.1038/s41586-020-1954-0.

15. Lee, Y., Warne, T., Nehmé, R., Pandey, S., Dwivedi-Agnihotri, H., Chaturvedi, M., Edwards, P.C., García-Nafría, J., Leslie, A.G.W., Shukla, A.K., et al. (2020). Molecular basis of β-arrestin coupling to formoterol-bound β1-adrenoceptor. Nature 583, 862–866. 10.1038/s41586-020-2419-1.

16. Bous, J., Fouillen, A., Orcel, H., Trapani, S., Cong, X., Fontanel, S., Saint-Paul, J., Lai-Kee-Him, J., Urbach, S., Sibille, N., et al. (2022). Structure of the vasopressin hormone–V2 receptor–β-arrestin1 ternary complex. Science Advances. 10.1126/sciadv.abo7761.

17. Yin, W., Li, Z., Jin, M., Yin, Y.-L., de Waal, P.W., Pal, K., Yin, Y., Gao, X., He, Y., Gao, J., et al. (2019). A complex structure of arrestin-2 bound to a G protein-coupled receptor. Cell Res 29, 971–983. 10.1038/s41422-019-0256-2.

18. Huang, W., Masureel, M., Qu, Q., Janetzko, J., Inoue, A., Kato, H.E., Robertson, M.J., Nguyen, K.C., Glenn, J.S., Skiniotis, G., et al. (2020). Structure of the neurotensin receptor 1 in complex with β-arrestin 1. Nature 579, 303–308. 10.1038/s41586-020-1953-1.

19. He, Q.-T., Xiao, P., Huang, S.-M., Jia, Y.-L., Zhu, Z.-L., Lin, J.-Y., Yang, F., Tao, X.-N., Zhao, R.-J., Gao, F.-Y., et al. (2021). Structural studies of phosphorylation-dependent interactions between the V2R receptor and arrestin-2. Nat Commun 12, 2396. 10.1038/s41467-021-22731-x.

20. Shukla, A.K., Manglik, A., Kruse, A.C., Xiao, K., Reis, R.I., Tseng, W.-C., Staus, D.P., Hilger, D., Uysal, S., Huang, L.-Y., et al. (2013). Structure of active β-arrestin-1 bound to a G-protein-coupled receptor phosphopeptide. Nature 497, 137–141. 10.1038/nature12120.

21. Min, K., Yoon, H.-J., Park, J.Y., Baidya, M., Dwivedi-Agnihotri, H., Maharana, J., Chaturvedi, M., Chung, K.Y., Shukla, A.K., and Lee, H.H. (2020). Crystal Structure of β-Arrestin 2 in Complex with CXCR7 Phosphopeptide. Structure 28, 1014–1023.e4. 10.1016/j.str.2020.06.002.

22. Mayer, D., Damberger, F.F., Samarasimhareddy, M., Feldmueller, M., Vuckovic, Z., Flock, T., Bauer, B., Mutt, E., Zosel, F., Allain, F.H.T., et al. (2019). Distinct G protein-coupled receptor phosphorylation motifs modulate arrestin affinity and activation and global conformation. Nat Commun 10, 1261. 10.1038/s41467-019-09204-y.

23. Isaikina, P., Tsai, C.-J., Dietz, N., Pamula, F., Grahl, A., Goldie, K.N., Guixà-González, R., Branco, C., Paolini-Bertrand, M., Calo, N., et al. (2021). Structural basis of the activation of the CC chemokine receptor 5 by a chemokine agonist. Science Advances 7, eabg8685. 10.1126/sciadv.abg8685.

24. Scurci, I., Martins, E., and Hartley, O. (2018). CCR5: Established paradigms and new frontiers for a ‘celebrity’ chemokine receptor. Cytokine 109, 81–93. 10.1016/j.cyto.2018.02.018.

25. Alkhatib, G. (2009). The biology of CCR5 and CXCR4: Current Opinion in HIV and AIDS 4, 96–103. 10.1097/COH.0b013e328324bbec.

26. Jiao, X., Nawab, O., Patel, T., Kossenkov, A.V., Halama, N., Jaeger, D., and Pestell, R.G. (2019). Recent Advances Targeting CCR5 for Cancer and Its Role in Immuno-Oncology. Cancer Research 79, 4801–4807. 10.1158/0008-5472.CAN-19-1167.

27. Kraus, S., Kolman, T., Yeung, A., and Deming, D. (2021). Chemokine Receptor Antagonists: Role in Oncology. Curr Oncol Rep 23, 131. 10.1007/s11912-021-01117-8.

28. Chua, R.L., Lukassen, S., Trump, S., Hennig, B.P., Wendisch, D., Pott, F., Debnath, O., Thürmann, L., Kurth, F., Völker, M.T., et al. (2020). COVID-19 severity correlates with airway epithelium-immune cell interactions identified by single-cell analysis. Nat Biotechnol 38, 970–979. 10.1038/s41587-020-0602-4.

29. Sharma, D., and Parameswaran, N. (2015). Multifaceted role of β-arrestins in inflammation and disease. Genes Immun 16, 499–513. 10.1038/gene.2015.37.

30. Martins, E., Brodier, H., Rossitto-Borlat, I., Ilgaz, I., Villard, M., and Hartley, O. (2020). Arrestin Recruitment to C-C Chemokine Receptor 5: Potent C-C Chemokine Ligand 5 Analogs Reveal Differences in Dependence on Receptor Phosphorylation and Isoform-Specific Recruitment Bias. Mol Pharmacol 98, 599–611. 10.1124/molpharm.120.000036.

31. Oppermann, M., Mack, M., Proudfoot, A.E.I., and Olbrich, H. (1999). Differential Effects of CC Chemokines on CC Chemokine Receptor 5 (CCR5) Phosphorylation and Identification of Phosphorylation Sites on the CCR5 Carboxyl Terminus*. Journal of Biological Chemistry 274, 8875–8885. 10.1074/jbc.274.13.8875.

32. Aramori, I., Ferguson, S.S., Bieniasz, P.D., Zhang, J., Cullen, B., and Cullen, M.G. (1997). Molecular mechanism of desensitization of the chemokine receptor CCR-5: receptor signaling and internalization are dissociable from its role as an HIV-1 co-receptor. EMBO J 16, 4606–4616. 10.1093/emboj/16.15.4606.

33. Kleibeuker, W., Jurado-Pueyo, M., Murga, C., Eijkelkamp, N., Mayor Jr, F., Heijnen, C.J., and Kavelaars, A. (2008). Physiological changes in GRK2 regulate CCL2-induced signaling to ERK1/2 and Akt but not to MEK1/2 and calcium. Journal of Neurochemistry 104, 979–992. 10.1111/j.1471-4159.2007.05023.x.

34. Olbrich, H., Proudfoot, A.E.I., and Oppermann, M. (1999). Chemokine-induced phosphorylation of CC chemokine receptor 5 (CCR5). Journal of Leukocyte Biology 65, 281–285. 10.1002/jlb.65.3.281.

35. Vroon, A., Heijnen, C.J., and Kavelaars, A. (2006). GRKs and arrestins: regulators of migration and inflammation. Journal of Leukocyte Biology 80, 1214–1221. 10.1189/jlb.0606373.

36. Xiao, K., Shenoy, S.K., Nobles, K., and Lefkowitz, R.J. (2004). Activation-dependent Conformational Changes in β-Arrestin 2*. Journal of Biological Chemistry 279, 55744–55753. 10.1074/jbc.M409785200.

37. Han, M., Gurevich, V.V., Vishnivetskiy, S.A., Sigler, P.B., and Schubert, C. (2001). Crystal Structure of β-Arrestin at 1.9 Å: Possible Mechanism of Receptor Binding and Membrane Translocation. Structure 9, 869–880. 10.1016/S0969-2126(01)00644-X.

38. Kang, D.S., Kern, R.C., Puthenveedu, M.A., von Zastrow, M., Williams, J.C., and Benovic, J.L. (2009). Structure of an Arrestin2-Clathrin Complex Reveals a Novel Clathrin Binding Domain That Modulates Receptor Trafficking. J Biol Chem 284, 29860–29872. 10.1074/jbc.M109.023366.

39. Baidya, M., Kumari, P., Dwivedi-Agnihotri, H., Pandey, S., Sokrat, B., Sposini, S., Chaturvedi, M., Srivastava, A., Roy, D., Hanyaloglu, A.C., et al. (2020). Genetically encoded intrabody sensors report the interaction and trafficking of β-arrestin 1 upon activation of G-protein–coupled receptors. Journal of Biological Chemistry 295, 10153–10167. 10.1074/jbc.RA120.013470.

40. Baidya, M., Kumari, P., Dwivedi-Agnihotri, H., Pandey, S., Chaturvedi, M., Stepniewski, T.M., Kawakami, K., Cao, Y., Laporte, S.A., Selent, J., et al. (2020). Key phosphorylation sites in GPCR s orchestrate the contribution of β-Arrestin 1 in ERK 1/2 activation. EMBO Rep 21. 10.15252/embr.201949886.

41. Dwivedi-Agnihotri, H., Chaturvedi, M., Baidya, M., Stepniewski, T.M., Pandey, S., Maharana, J., Srivastava, A., Caengprasath, N., Hanyaloglu, A.C., Selent, J., et al. (2020). Distinct phosphorylation sites in a prototypical GPCR differently orchestrate β-arrestin interaction, trafficking, and signaling. Sci. Adv. 6, eabb8368. 10.1126/sciadv.abb8368.

42. Shiraishi, Y., Kofuku, Y., Ueda, T., Pandey, S., Dwivedi-Agnihotri, H., Shukla, A.K., and Shimada, I. (2021). Biphasic activation of β-arrestin 1 upon interaction with a GPCR revealed by methyl-TROSY NMR. Nat Commun 12, 7158. 10.1038/s41467-021-27482-3.

43. Isberg, V., de Graaf, C., Bortolato, A., Cherezov, V., Katritch, V., Marshall, F.H., Mordalski, S., Pin, J.-P., Stevens, R.C., Vriend, G., et al. (2015). Generic GPCR residue numbers – aligning topology maps while minding the gaps. Trends in Pharmacological Sciences 36, 22–31. 10.1016/j.tips.2014.11.001.

44. Scott, M.G.H., Le Rouzic, E., Périanin, A., Pierotti, V., Enslen, H., Benichou, S., Marullo, S., and Benmerah, (2002). Differential Nucleocytoplasmic Shuttling of β-Arrestins: CHARACTERIZATION OF A LEUCINE-RICH NUCLEAR EXPORT SIGNAL IN β-ARRESTIN2*. Journal of Biological Chemistry 277, 37693–37701. 10.1074/jbc.M207552200.

45. Wang, P., Wu, Y., Ge, X., Ma, L., and Pei, G. (2003). Subcellular Localization of β-Arrestins Is Determined by Their Intact N Domain and the Nuclear Export Signal at the C Terminus*. Journal of Biological Chemistry 278, 11648–11653. 10.1074/jbc.M208109200.

46. Oakley, R.H., Laporte, S.A., Holt, J.A., Caron, M.G., and Barak, L.S. (2000). Differential Affinities of Visual Arrestin, βArrestin1, and βArrestin2 for G Protein-coupled Receptors Delineate Two Major Classes of Receptors*. Journal of Biological Chemistry 275, 17201–17210. 10.1074/jbc.M910348199.

47. Asher, W.B., Terry, D.S., Gregorio, G.G.A., Kahsai, A.W., Borgia, A., Xie, B., Modak, A., Zhu, Y., Jang, W., Govindaraju, A., et al. (2022). GPCR-mediated β-arrestin activation deconvoluted with single-molecule precision. Cell 185, 1661–1675.e16. 10.1016/j.cell.2022.03.042.

48. Isaikina, P., Tsai, C.-J., Petrovic, I., Rogowski, M., Dürr, A.M., and Grzesiek, S. (2022). Preparation of a stable CCL5·CCR5·Gi signaling complex for Cryo-EM analysis. Methods Cell Biol 169, 115–141. 10.1016/bs.mcb.2022.03.001.

49. Inglese, J., Koch, W.J., Caron, M.G., and Lefkowitz, R.J. (1992). Isoprenylation in regulation of signal transduction by G-protein-coupled receptor kinases. Nature 359, 147–150. 10.1038/359147a0.

50. Delaglio, F., Grzesiek, S., Vuister, G.W., Zhu, G., Pfeifer, J., and Bax, A. (1995). NMRPipe: A multidimensional spectral processing system based on UNIX pipes. J Biomol NMR 6, 277–293. 10.1007/BF00197809.

51. Lee, W., Tonelli, M., and Markley, J.L. (2015). NMRFAM-SPARKY: enhanced software for biomolecular NMR spectroscopy. Bioinformatics 31, 1325–1327. 10.1093/bioinformatics/btu830.

52. Kabsch, W. (2010). XDS. Acta Crystallogr. D Biol. Crystallogr. 66, 125–132. 10.1107/S0907444909047337.

53. Winn, M.D., Ballard, C.C., Cowtan, K.D., Dodson, E.J., Emsley, P., Evans, P.R., Keegan, R.M., Krissinel, E.B., Leslie, A.G.W., McCoy, A., et al. (2011). Overview of the CCP4 suite and current developments. Acta Cryst D 67, 235–242. 10.1107/S0907444910045749.

54. Bricogne, G., Blanc, E., Brandl, M., Flensburg, C., Keller, P., Paciorek, W., Roversi, P., Sharff, A., Smart, O.S., and Vonrhein, C. (2011). BUSTER version 2.10. 0. Global Phasing Ltd, Cambridge, UK.

55. Adams, P.D., Grosse-Kunstleve, R.W., Hung, L.-W., Ioerger, T.R., McCoy, A.J., Moriarty, N.W., Read, R.J., Sacchettini, J.C., Sauter, N.K., and Terwilliger, T.C. (2002). PHENIX: building new software for automated crystallographic structure determination. Acta Cryst D 58, 1948–1954. 10.1107/S0907444902016657.

56. Inoue, A., Raimondi, F., Kadji, F.M.N., Singh, G., Kishi, T., Uwamizu, A., Ono, Y., Shinjo, Y., Ishida, S., Arang, N., et al. (2019). Illuminating G-Protein-Coupling Selectivity of GPCRs. Cell 177, 1933–1947.e25. 10.1016/j.cell.2019.04.044.

57. Pandey, S., Roy, D., and Shukla, A.K. (2019). Chapter 8 - Measuring surface expression and endocytosis of GPCRs using whole-cell ELISA. In Methods in Cell Biology G Protein-Coupled Receptors, Part B., A. K. Shukla, ed. (Academic Press), pp. 131–140. 10.1016/bs.mcb.2018.09.014.

