## Supporting Information for "A key GPCR phosphorylation motif discovered in arrestin2•CCR5 phosphopeptide complexes"

*Address correspondence to:
Stephan Grzesiek
Focal Area Structural Biology and Biophysics, Biozentrum
University of Basel, CH-4056 Basel, Switzerland
Phone: ++41 61 267 2100


Arun K. Shukla
Department of Biological Sciences and Bioengineering
Indian Institute of Technology, Kanpur 208016, India
Phone:


Polina Isaikina
Focal Area Structural Biology and Biophysics, Biozentrum
University of Basel, CH-4056 Basel, Switzerland
Phone:


### Supplementary Tables

**Table S1.** Statistics on diffraction data and refinement of human arrestin2 (apo) and CCR5pp•arrestin2•Fab30 complexes.

| | 4P•arrestin2•Fab30 | 6P•arrestin2•Fab30 | human arrestin2 |
|---|---|---|---|
| PDB Identifier | 8AS2 | 8AS3 | 8AS4 |
| Wavelength (Å) | 1.00000 | 0.99999 | 0.999999 |
| Resolution range (Å) | 46.8 – 3.2 (3.31 – 3.2)* | 46.5- 3.50 (3.63 – 3.50)* | 44.75 – 2.30 (2.38 – 2.30)* |
| Space group | I 21 21 21 | I 21 21 21 | P 1 21 1 |
| Unit cell dimensions (Å) | 116.8 122.6 145.9 | 116.3 121.1 145.3 | 62.0 71.4 115.4 |
| α, β, γ (°) | 90.0 90 90.0 | 90.0 90.0 90.0 | 90.0 97.7 90.0 |
| Total reflections | 461,526 (42,112) | 178,306 (17,607) | 1,216,192 (119,044) |
| Unique reflections | 17,524 (1,702) | 13,152 (1,295) | 44,486 (4,409) |
| Multiplicity | 26.3 (26.8) | 13.6 (13.6) | 27.3 (27.0) |
| Completeness (%) | 99.9 (100) | 98.3 (90.3) | 99.8 (99.0) |
| Mean I/sigma(I) | 11.6 (1.1) | 12.2 (0.7) | 12.1 (1.6) |
| Wilson B-factor | 120.1 | 171.8 | 48.6 |
| R-merge (%) | 22.0 (399.2) | 14.4 (525.4) | 24.0 (246.1) |
| Rpim (%) | 4.4 (78.9) | 4.0 (146.5) | 7.0 (77.9) |
| CC1/2 | 0.998 (0.662) | 0.999 (0.317) | 0.999 (0.790) |
| Reflections used in Refinement | 17,439 (1,691) | 13,005 (1,171) | 44,388 (4,397) |
| R-work | 0.281 (0.428) | 0.283 (0.442) | 0.224 (0.361) |
| R-free | 0.317 (0.488) | 0.335 (0.454) | 0.252 (0.407) |
| Number of atoms | 6,096 | 6,067 | 5,892 |
| Protein residues | 788 | 783 | 325 |
| RMSD(bonds) (Å) | 0.003 | 0.004 | 0.011 |
| RMSD(angles) (°) | 0.68 | 0.75 | 1.66 |
| Ramachandran favored (%) | 94.0 | 96.0 | 96.5 |
| Ramachandran outliers (%) | 0 | 0 | 0 |
| Rotamer outliers (%) | 0 | 0 | 0.5 |
| Clash score | 2.30 | 4.87 | 3.78 |
| Average B-factor | 197.4 | 199.9 | 80.3 |

*The values in parentheses correspond to the highest resolution shell.

**Table S2**. GPCR classification based of the characteristics of agonist-dependent arrestin interactions.

| Receptor name | Class | Method | Reference |
|---|---|---|---|
| α1B-ADR | A | Functional assays | (Oakley et al. 2000) |
| β2-ADR | A | Functional assays | (Oakley et al. 2000) |
| μOR | A | Functional assays | (Oakley et al. 2000) |
| ETAR | A | Functional assays | (Oakley et al. 2000) |
| D1R | A | Functional assays | (Oakley et al. 2000) |
| D2R | A | Functional assays | (Asher et al. 2022) |
| 5HT2aR | A | Functional assays | (Schmid, Raehal, and Bohn 2008) |
| 5HT2bR | A | Functional and structural data | (Cao et al. 2022) |
| M1R | A | FRET | (Jung et al. 2017) |
| M2R | A | Functional and structural data on M2R-V2Rpp (indirect evidence*)* | (Staus et al. 2018; 2020) |
| SST3R | A | Functional assays | (Tulipano et al. 2004) |
| SST5R | A | Functional assays | (Tulipano et al. 2004) |
| CXCR3B | A | Functional assays | (Smith et al. 2017) |
| CXCR3A | B | Functional assays | (Smith et al. 2017) |
| CXCR4 | B | Functional assays | (Liebick et al. 2017) |
| CCR5 | B | Functional assays | (Liebick et al. 2017) |
| CCR7 | B | Functional assays | (Zidar et al. 2009) |
| ACKR2 | B | Functional assays | (Pandey et al. 2021) |
| ACKR3 | B | Functional assays and structural data | (Rajagopal et al. 2010; Sarma et al. 2022) |
| V2R | B | Functional assays and structural data | (Oakley et al. 2000) |
| NTR1 | B | Functional assays and structural data | (Oakley et al. 2000) |
| C5aR2 | B | Functional assays | (Pandey et al. 2021) |
| Rho | B | Biochemical and biophysical data | (Mayer et al. 2019) |
| AT1R | B | Functional assays | (Oakley et al. 2000) |
| OTR | B | Functional assays | (Oakley et al. 2000) |
| SST2R | B | Functional assays | (Tulipano et al. 2004) |
| TRH1R | B | Functional assays | (Oakley et al. 2000) |

### Supplementary Figures

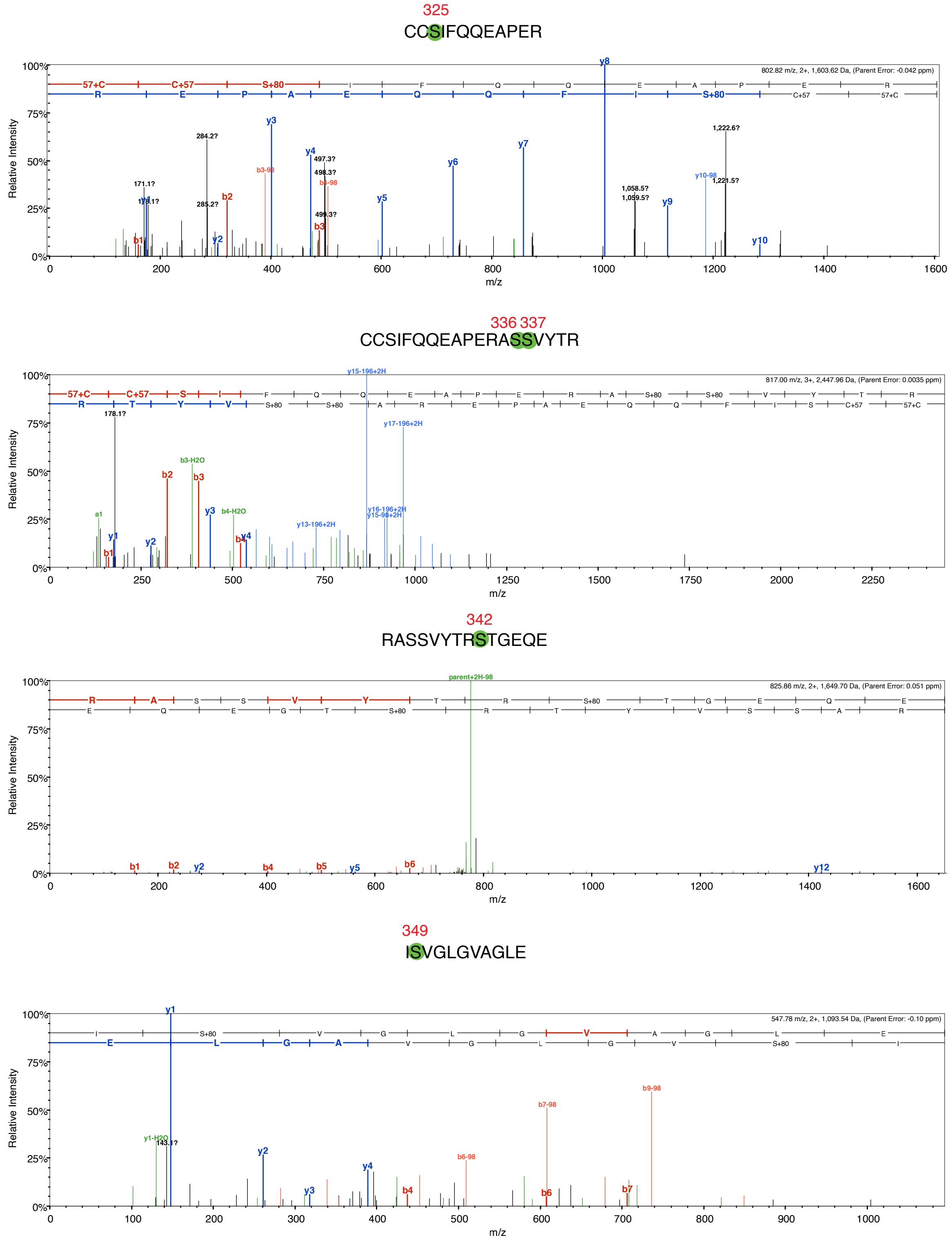


**Figure S1**. Mass spectra used for detection of CCR5 phosphorylation induced by GRK2. Representative spectra of the identified peptides bearing phosphorylated residues. Residues with higher occupancies are highlighted by green circles.

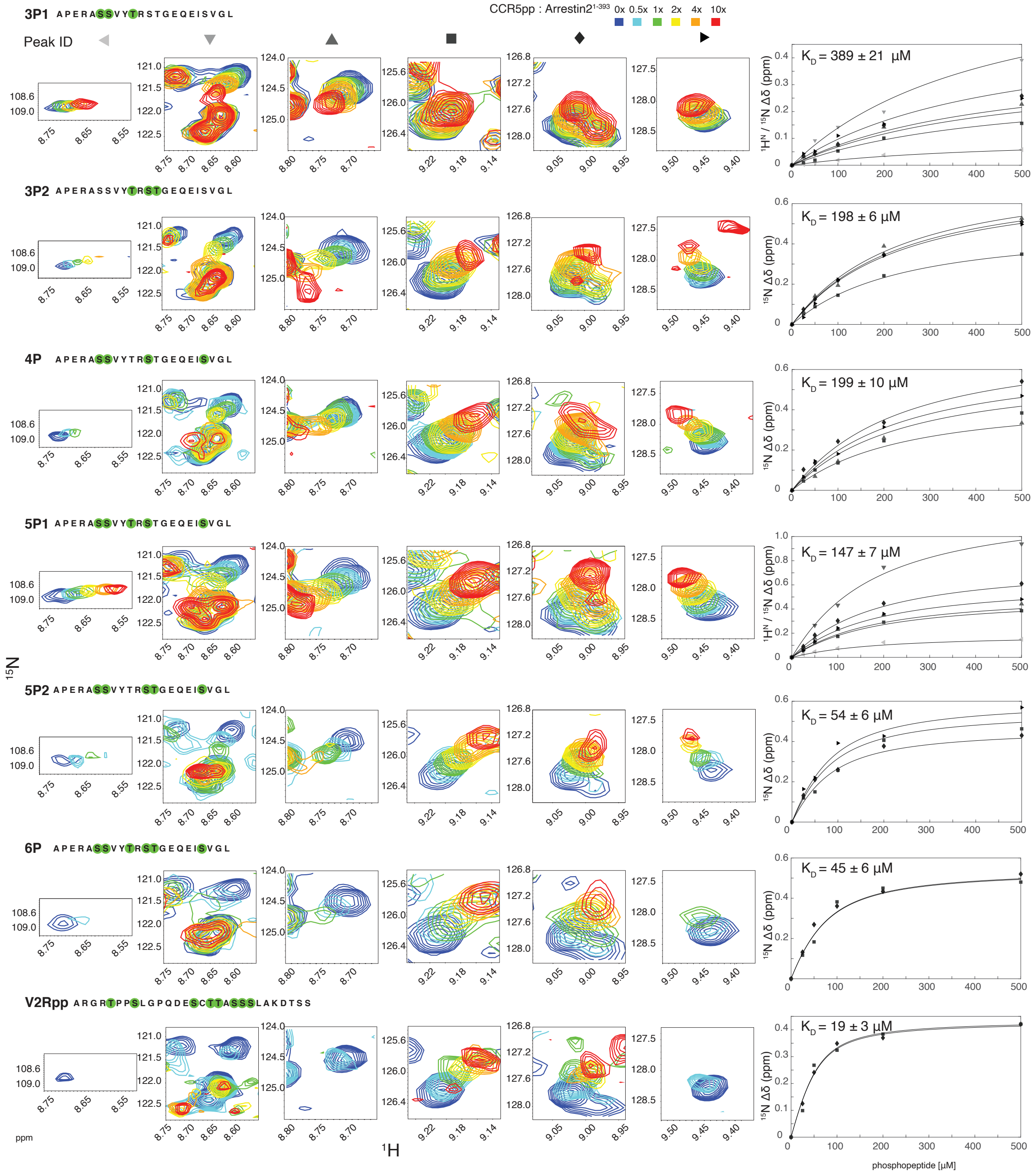


**Figure S2.** NMR titration of arrestin2$^{1-393}$ with CCR5 phosphopeptides. Left: small regions of $^{1}H$-$^{15}N$ TROSY spectra showing resonance shifts of four selected arrestin2$^{1-393}$ residues upon CCR5 phosphopeptide binding. The individual peptides are indicated at the top of each row. Right: detected chemical shift changes as a function of phosphopeptide concentration. Solid lines depict global non-linear least-square fits to the data points with respective dissociation constants (see Methods).

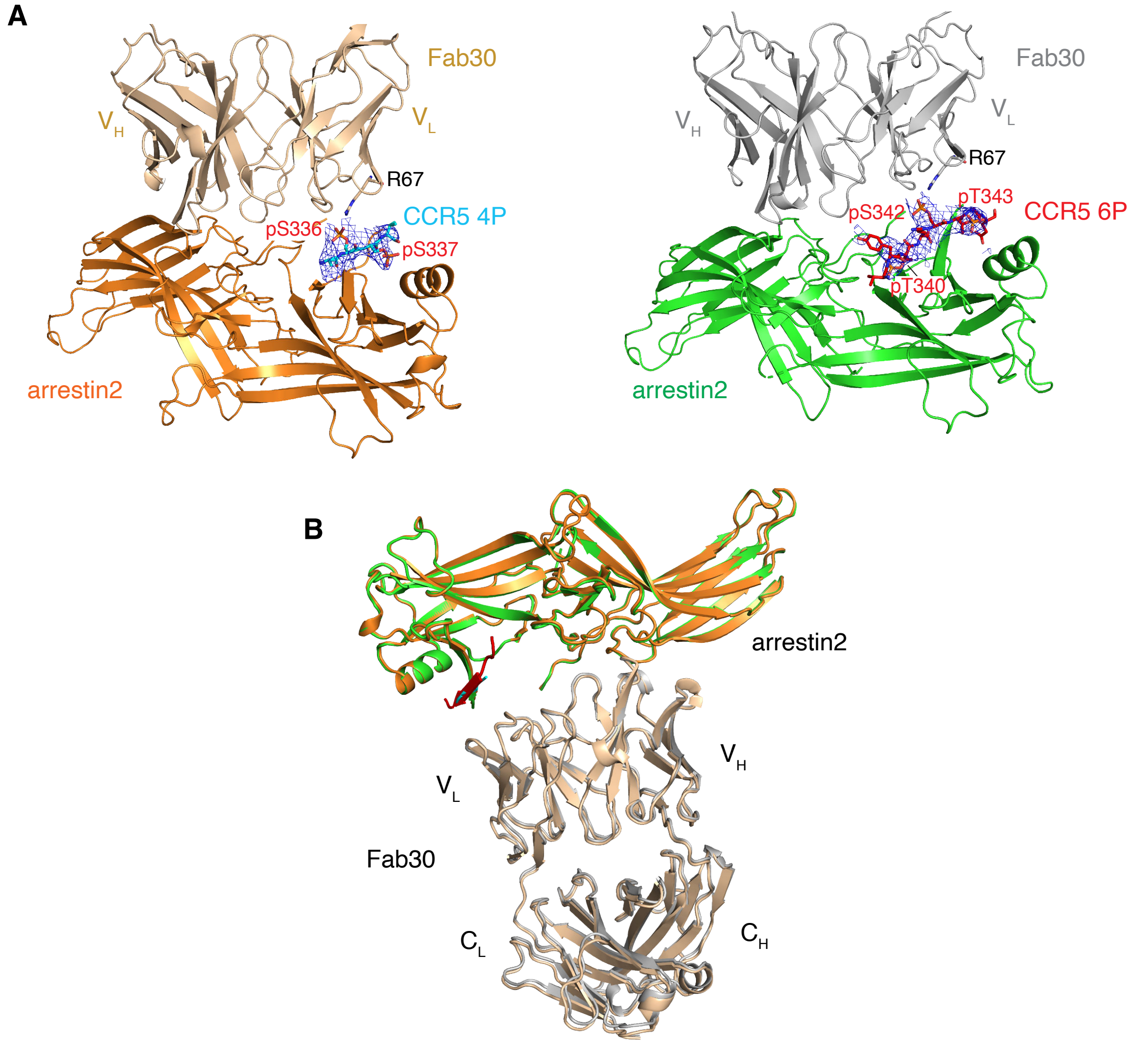


**Figure S3.** Phosphopeptide electron density and Fab30 coordination in arrestin2•CCR5pp crystal structures. (A) 2Fo-Fc electron density maps of the CCR5 4P (left) and 6P (right) phosphopeptides and contacts between the phosphopeptides and Fab30. The electron densities are contoured at 1.0 σ cut-off within 6 Å and displayed as a blue mesh. (B) Overall coordination of the 6P•arrestin2 and 4P•arrestin2 complexes with Fab30. Fab30 is colored in beige in the 4P complex and in light gray in the 6P complex. Only very minor conformation changes of Fab30 upon binding to the two CCR5 phosphopeptides are detectable.

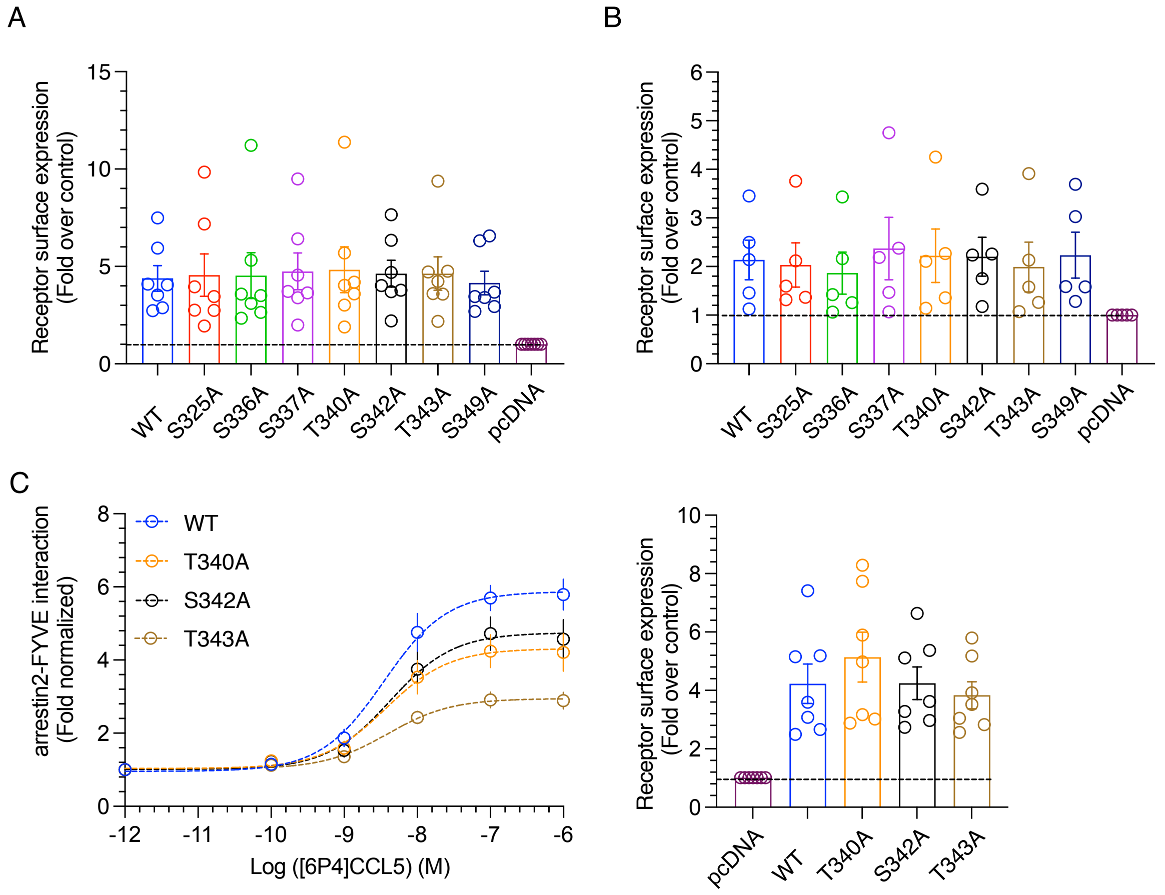


**Figure S4.** Arrestin2 trafficking assay and surface expression of the various CCR5 phospho-deficient mutants in different cellular assays. (A) Surface expression levels of wild-type CCR5 and all S/A and T/A CCR5 mutants used in the Ib30-based NanoBiT assay. (B) Surface expression levels of wild-type CCR5 and all S/A and T/A CCR5 mutants used in the arrestin2 recruitment NanoBiT assay. (C) Left: arrestin2 trafficking assay on wild-type CCR5 and selected CCR5 mutants (T340A, S342A and T343A) in response to [6P4]CCL5 stimulation. Right: surface expression of the CCR5 constructs used in this assay.

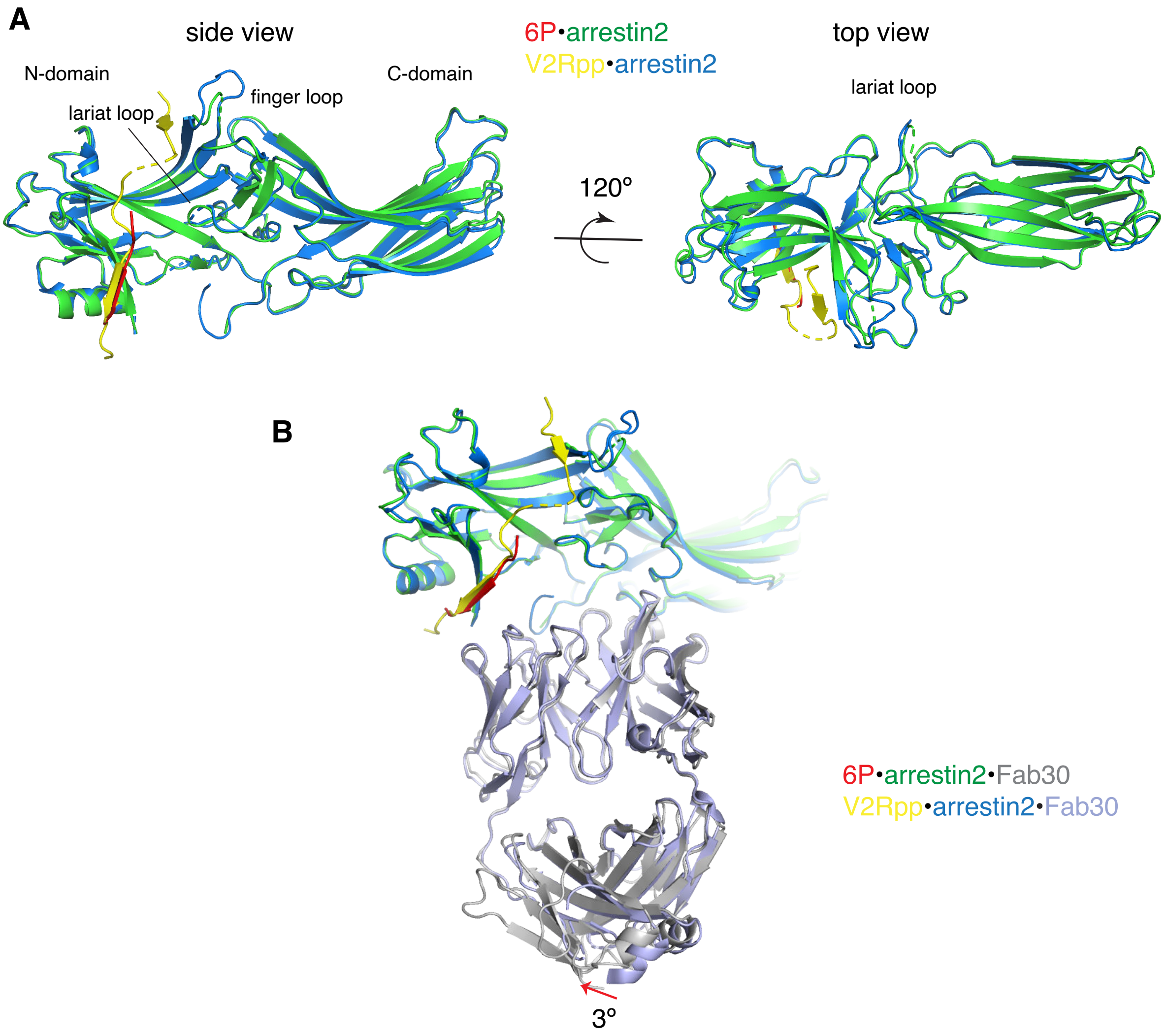


**Figure S5.** Overall structural comparison between 6P•arrestin2•Fab30 and V2Rpp•arrestin2•Fab30. (A) Superposition of 6P•arrestin2•Fab30 and V2Rpp•arrestin2•Fab30 structures in side and top views. The color code is indicated and identical to Figure 5A. (B) Overall Fab30 binding in both structures. The color code is indicated and follows panel A.

### Supplementary References


Asher, Wesley B., Daniel S. Terry, G. Glenn A. Gregorio, Alem W. Kahsai, Alessandro Borgia, Bing Xie, Arnab Modak, et al. 2022. "GPCR-Mediated β-Arrestin Activation Deconvoluted with Single-Molecule Precision." *Cell* 185 (10): 1661-1675.e16. https://doi.org/10.1016/j.cell.2022.03.042.

Cao, Can, Ximena Barros-Álvarez, Shicheng Zhang, Kuglae Kim, Marc A. Dämgen, Ouliana Panova, Carl-Mikael Suomivuori, et al. 2022. "Signaling Snapshots of a Serotonin Receptor Activated by the Prototypical Psychedelic LSD." *Neuron*, September. https://doi.org/10.1016/j.neuron.2022.08.006.

Jung, Seung-Ryoung, Christopher Kushmerick, Jong Bae Seo, Duk-Su Koh, and Bertil Hille. 2017. "Muscarinic Receptor Regulates Extracellular Signal Regulated Kinase by Two Modes of Arrestin Binding." *Proceedings of the National Academy of Sciences* 114 (28): E5579–88. https://doi.org/10.1073/pnas.1700331114.

Liebick, Marcel, Sarah Henze, Viola Vogt, and Martin Oppermann. 2017. "Functional Consequences of Chemically-Induced β-Arrestin Binding to Chemokine Receptors CXCR4 and CCR5 in the Absence of Ligand Stimulation." *Cellular Signalling* 38 (October): 201–11. https://doi.org/10.1016/j.cellsig.2017.07.010.

Mayer, Daniel, Fred F. Damberger, Mamidi Samarasimhareddy, Miki Feldmueller, Ziva Vuckovic, Tilman Flock, Brian Bauer, et al. 2019. "Distinct G Protein-Coupled Receptor Phosphorylation Motifs Modulate Arrestin Affinity and Activation and Global Conformation." *Nature Communications* 10 (1): 1261. https://doi.org/10.1038/s41467-019-09204-y.

Oakley, Robert H., Stéphane A. Laporte, Jason A. Holt, Marc G. Caron, and Larry S. Barak. 2000. "Differential Affinities of Visual Arrestin, BArrestin1, and BArrestin2 for G Protein-Coupled Receptors Delineate Two Major Classes of Receptors*." *Journal of Biological Chemistry* 275 (22): 17201–10. https://doi.org/10.1074/jbc.M910348199.

Pandey, Shubhi, Punita Kumari, Mithu Baidya, Ryoji Kise, Yubo Cao, Hemlata Dwivedi-Agnihotri, Ramanuj Banerjee, et al. 2021. "Intrinsic Bias at Non-Canonical, β-Arrestin-Coupled Seven Transmembrane Receptors." *Molecular Cell* 81 (22): 4605-4621.e11. https://doi.org/10.1016/j.molcel.2021.09.007.

Rajagopal, Sudarshan, Jihee Kim, Seungkirl Ahn, Stewart Craig, Christopher M. Lam, Norma P. Gerard, Craig Gerard, and Robert J. Lefkowitz. 2010. "β-Arrestin- but Not G Protein-Mediated Signaling by the 'Decoy' Receptor CXCR7." *Proceedings of the National Academy of Sciences* 107 (2): 628–32. https://doi.org/10.1073/pnas.0912852107.

Sarma, Parishmita, Hye-Jin Yoon, Carlo Marion C. Carino, Deeksha S, Ramanuj Banerjee, Yaejin Yun, Jeongsek Ji, et al. 2022. "Molecular Insights into Intrinsic Transducer-Coupling Bias in the CXCR4-CXCR7 System." *BioRxiv*, June, 2022.06.06.494935. https://doi.org/10.1101/2022.06.06.494935.

Schmid, Cullen L., Kirsten M. Raehal, and Laura M. Bohn. 2008. "Agonist-Directed Signaling of the Serotonin 2A Receptor Depends on β-Arrestin-2 Interactions *in Vivo*." *Proceedings of the National Academy of Sciences* 105 (3): 1079–84. https://doi.org/10.1073/pnas.0708862105.

Smith, Jeffrey S., Priya Alagesan, Nimit K. Desai, Thomas F. Pack, Jiao-Hui Wu, Asuka Inoue, Neil J. Freedman, and Sudarshan Rajagopal. 2017. "C-X-C Motif Chemokine Receptor 3 Splice Variants Differentially Activate Beta-Arrestins to Regulate Downstream Signaling Pathways." *Molecular Pharmacology* 92 (2): 136–50. https://doi.org/10.1124/mol.117.108522.

Staus, Dean P., Hongli Hu, Michael J. Robertson, Alissa L. W. Kleinhenz, Laura M. Wingler, William D. Capel, Naomi R. Latorraca, Robert J. Lefkowitz, and Georgios Skiniotis. 2020. "Structure of the M2 Muscarinic Receptor–β-Arrestin Complex in a Lipid Nanodisc." *Nature* 579 (7798): 297–302. https://doi.org/10.1038/s41586-020-1954-0.

Staus, Dean P., Laura M. Wingler, Minjung Choi, Biswaranjan Pani, Aashish Manglik, Andrew C. Kruse, and Robert J. Lefkowitz. 2018. "Sortase Ligation Enables Homogeneous GPCR Phosphorylation to Reveal Diversity in β-Arrestin Coupling." *Proceedings of the National Academy of Sciences* 115 (15): 3834–39. https://doi.org/10.1073/pnas.1722336115.

Tulipano, Giovanni, Ralf Stumm, Manuela Pfeiffer, Hans-Jürgen Kreienkamp, Volker Höllt, and Stefan Schulz. 2004. "Differential β-Arrestin Trafficking and Endosomal Sorting of Somatostatin Receptor Subtypes*." *Journal of Biological Chemistry* 279 (20): 21374–82. https://doi.org/10.1074/jbc.M313522200.

Zidar, David A., Jonathan D. Violin, Erin J. Whalen, and Robert J. Lefkowitz. 2009. “Selective Engagement of G Protein Coupled Receptor Kinases (GRKs) Encodes Distinct Functions of Biased Ligands.” *Proceedings of the National Academy of Sciences* 106 (24): 9649–54. https://doi.org/10.1073/pnas.0904361106.
